# An immunophenotype-coupled transcriptomic atlas of mouse hematopoietic progenitors

**DOI:** 10.64898/2026.09.08.750044

**Authors:** Mahamudul Hasan Bhuyan, Sumanta Barman, Shafaqat Ali, Felix Steinbeck, Tobias Lautwein, Norbert Goebels, Stefanie Scheu

## Abstract

Decoding the bone marrow (BM) hematopoietic progenitor compartment is central to understanding the development of mature immune cells in health and disease. Single-cell multimodal approaches are transforming the analysis of this heterogeneous compartment by simultaneously capturing transcriptomic and proteomic information. Still, integrated studies covering the full spectrum of murine hematopoietic stem and progenitor cells (HSPCs) remain scarce. Here, we report a Cellular Indexing of Transcriptomes and Epitopes by sequencing (CITE-seq) based atlas of 9,281 low-frequency mouse BM HSPCs, profiled using a 123 antibody-derived tag (ADT) panel coupled with genome-wide single-cell transcriptomes. Multimodal analysis resolved 28 progenitor stages across eight blood lineages, each defined by distinctive surface marker and transcription factor (TF) profiles, and provided novel marker combinations to dissect transcriptionally closely related progenitor stages. Classical mouse progenitor surface markers, such as CD117, CD34, and CD115, were detected in accordance with well-established lineage relationships, whereas CD43 and CD48 showed, unexpectedly, broader expression patterns. Neglected surface molecules such as CD27, CD36, CD106, and PIR-A/B are nominated as novel stage-specific markers. ADT versus transcriptome comparison revealed transcription-translation inconsistencies for markers, including CD54 and CD68. A CITE-seq-guided gating strategy translates these population definitions into FACS-compatible panels for prospective isolation and functional interrogation of previously unresolved progenitor stages. Comprehensive TF network analysis delineated lineage- and stage-specific transcriptional programs. This contributes to clarifying unresolved ontogenies and functions, including the lympho-myeloid dual origin of pDCs or rapid neutrophil turnover. This new and comprehensive resource enables exploration of mouse hematopoietic progenitor states driving discoveries in hematology and immunology.

**Key Points:** • An immunophenotypic and transcription factor map of murine bone marrow hematopoietic progenitor cells is presented.

• Surface marker vs transcriptome profiles enable better lineage identification, including pDC and neutrophil progenitors.

## 2. Introduction

Hematopoiesis is a dynamic process in which hematopoietic stem cells (HSCs) at the apex of the hierarchy both self-renew and give rise to progenitors. These progenitors pass through stages of fate decision, committing to the erythroid, myeloid, or lymphoid lineages.^1–5^ The ability to unequivocally identify and purify progenitors at specific differentiation stages is fundamental to improve leukemia characterization, BM transplantation, targeted therapies, and regenerative medicine strategies.

The progenitor compartment of the BM is composed of highly plastic cells with extensively overlapping surface markers, transcripts, and developmental trajectories, making them difficult to classify precisely. Specifically, the coherence between function and fate across many progenitor stages has been questioned by multiple studies, making it increasingly challenging to refine the hematopoietic lineage hierarchy.^6–9^

Cytometry-based analyses, such as FACS (fluorescence-assisted cell sorting), CyTOF (cytometry by time-of-flight), and spectral flow provide protein-level information. Despite the possibility to measure an increasing number of parameters, these methods remain hypothesis-based, relying on predefined cell-surface markers to distinguish specific progenitors, precluding the unbiased discovery of novel, possibly more precise and unique marker patterns.^4,10–12^

On the other hand, high-resolution single-cell RNA sequencing technologies now allow systematic characterization of TF landscapes, providing novel insights into hematopoietic stem and progenitor cell (HSPC) heterogeneity and dynamics during fate commitment and unbiased discovery of novel progenitor populations.^13–15^ However, these methods exclusively measure RNA, which does not always correlate directly with protein abundance, which is especially important for understanding cell function.^16,17^ Such methods also preclude isolation of live cells from the prospective cell clusters, which is essential for their functional evaluation and clinical translation.^18–21^

Collectively, findings generated by these methods underscore a gap between single-cell sequencing-based cell type classification and data from multiplexed FACS assays.^22,23^ The recently introduced technology of high-resolution immunophenotypic and transcriptional state-coupled multi-omics, cellular indexing of transcriptomes and epitopes by sequencing (CITE-seq), integrates surface markers with fate-defining TFs, thereby allowing characterization, isolation, and functional analysis of hematopoietic stem cells, progenitor populations, and their transitional compartments.^24^ This approach implements theoretically unlimited multiplexing of antibody-derived tags (ADTs), a powerful framework for the identification of novel markers and the definition of previously unrecognized populations with unique marker constellations beyond transcriptional states.^25^ In recent years, single-cell proteogenomic approaches, such as CITE-seq, have increasingly been applied to study the hematopoietic system. Although the majority of these studies have focused on human hematopoiesis, several mouse datasets have also been generated. However, the hematopoietic progenitor compartment remains underrepresented across these studies due to the lineage-specific enrichment strategies used. Moreover, the limited number of CITE-seq antibodies included in most studies has hindered the identification of novel surface marker combinations within the hematopoietic progenitor compartment.^25–29^

In this study, we set out to discover and profile rare HSPCs using CITE-seq in the mouse as the primary model organism for functional hematopoiesis research. Comparative analysis of molecular signatures during hematopoietic development allowed us to delineate progenitor stage-specific core signatures, derived from both TFs and surface epitopes. CITE-seq enabled us to directly compare immunophenotypes and transcriptomes for better projection of the diversity and complexity of the examined early hematopoietic stages. This work provides a foundational reference and overarching framework for further functional investigation on hematopoietic progenitors at single-cell resolution. We further provide a comprehensive single-cell gene and surface marker expression atlas as a publicly available online resource for navigating HSC-to-progenitor transitions, lineage bifurcations, as well as lineage-defining transcriptional programs.

## 3. Methods

### Preparation of BM and cell sorting

Lin^-^ HSPCs were enriched by negative FACS sorting against the surface markers CD3, CD4, CD8a, CD11b, CD11c, CD19, B220, MHC-II, Ly-6G, NK1.1, and Ter119 (supplemental Table 1) from BM cells extracted from six 10-weeks-old C57BL/6 mice.^30^ Detailed gating strategy is shown in supplemental Figure 1.

### Sample preparation for CITE-seq

Sorted cells from six individual animals were tagged using TotalSeq^TM^-A anti-mouse Hashtag antibodies 1-6 (supplemental Table 2), washed 3x, and pooled. CITE-seq staining was performed with the TotalSeq^TM^-A Mouse Universal Cocktail-v1.0 (119 antibodies and 9 isotypes) with spiked-in pre-titrated amounts of four additional CITE-seq antibodies against CD117, CD34, CD135, and CD16/32 (supplemental Tables 3 and 4).

### Single-cell CITE-seq library generation and sequencing

20,000 cells were used as input for the single-cell droplet library generation on the 10X Chromium Controller system. Sequencing was carried out with a mean sequencing depth of 50,000 reads/cell for the Gene Expression library, ∼15,000 reads/cell for the ADT library, and ∼5,000 reads/cell for the Cell Hashing library.

### Data processing and bioinformatic analysis

Raw sequencing data was preprocessed using CellRanger software (v7.0) aligning it to the mm10 genome. Seurat^31^ (v5.2.1) package was used for quality control, integration, and visualization of the data. An integrated weighted nearest neighbor (wnn) UMAP^32^ embedding was generated using the FindMultiModalNeighbors function. Pseudotime trajectory along differentiation was analyzed with Monocle3^33^ (v1.4.27). A detailed description of sequencing, data processing, and analysis is given in the supplemental methods.

## 4. Results

### CITE-seq profiling of BM progenitors defines 28 manually curated clusters attributed to eight distinct hematopoietic lineages

To capture a comprehensive spectrum of hematopoietic progenitors from mouse BM for integrated single-cell analysis of the surface marker phenotype in conjunction with the transcriptome using CITE-Seq, we enriched lin^-^ progenitor cells through an unbiased gating strategy, avoiding selective targeting of any progenitor type (Figure 1A; supplemental Figure 1). The resulting dataset included 9,281 individual cells with a median detection of 4,477 genes per cell and 123 ADT-derived surface markers.

**Figure 1.**
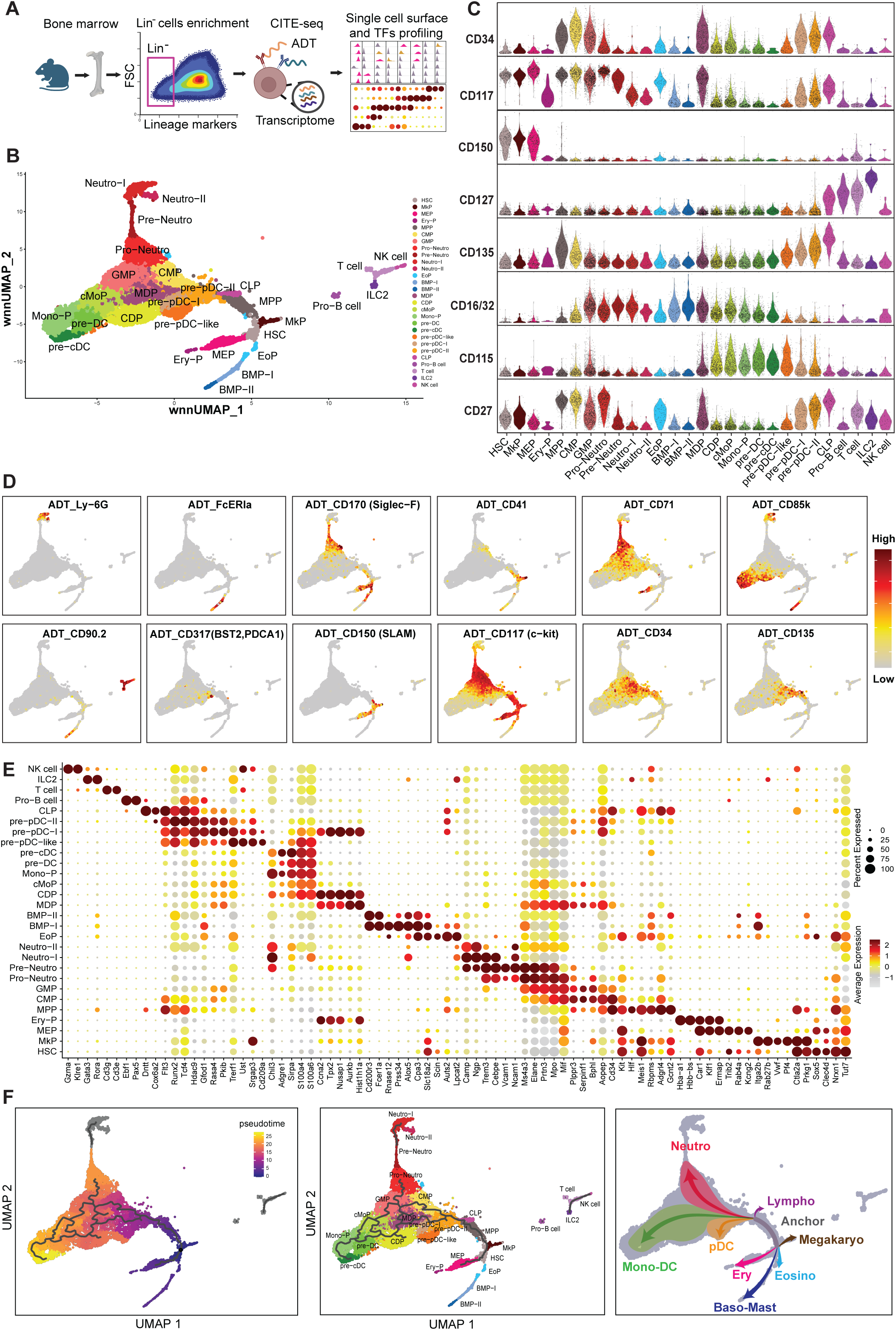
An integrative single-cell transcriptional and surface marker profile of mouse BM progenitor cell states. (A) Schematic overview of bone marrow preparation, lin^-^ cell enrichment, and CITE-seq workflow. (B) wnnUMAP projection of single-cell proteo-genomics data of mouse lin^-^ BM progenitor cells (n=9,281 single cells, 123 surface markers). A total of 28 distinct cellular states were defined by clustering and annotated based on cell-surface markers and signature transcripts. Cells are colored by clusters. (C) Violin plots depicting the cluster-specific expression of eight well-established hallmark surface markers for the definition of BM progenitors. (D) wnnUMAP overlays illustrating protein expression patterns of bone marrow lineage defining surface markers. The portion of the wnnUMAP enriched with lineage/progenitor specific surface proteins represent the corresponding lineage branch. The scale bar applies to the scaled expression level shown in the panel. (E) Bead plot illustrating the expression of specific lineage defining marker transcripts across 28 cellular states. Mean expression level is represented as colored-scale of relative intensities and the size of the circle represents the fraction of cells within that cluster having non-zero expression values. (F) Monocle3 inferred wnnUMAP embedded pseudotime ordering (left) and trajectories (middle) of cellular states. A lineage branching model (right) is illustrated based on pseudotime trajectory and cell type identity markers. Abbreviations: HSC, hematopoietic stem cell; MkP, megakaryocyte progenitor; MEP, megaerythoid progenitor; Ery-P, erythroid progenitor; MPP, multipotent progenitor; CMP, common myeloid progenitor; GMP, granulocyte monocyte progenitor; Neutro, neutrophil; EoP, eosinophil progenitor; BMP, basophil/mast cell progenitor; MDP, monocyte dendritic cell progenitor; CDP, common dendritic cell progenitor; cMoP, common monocyte progenitor; Mono-P, monocyte progenitor; DC, dendritic cell; cDC, classical dendritic cell; pDC, plasmacytoid dendritic cell; CLP, common lymphoid progenitor.

We generated an unsupervised wnnUMAP^32^ plot based on a weighted combination of RNA and protein expression of surface markers (Figure 1B). Clusters were annotated by referring to well-established progenitor- (Figure 1C) and lineage-defining (Figure 1D) surface markers as well as lineage-associated transcripts (Figure 1E). This resulted in the definition of 28 manually curated clusters that could be attributed to eight unique blood lineages.

The surface markers TCRβ/CD3/CD5/CD8α/CD8β, CD19/B220, NK1.1/Ly49D, TER119, and Ly-6G allowed clear identification of the remnants of mature cell populations in our samples, such as T, B, NK, and erythroid cells, and neutrophils, respectively (supplemental Figure 2). CD150/CD117/CD34/CD135 outlined stem-like and multilineage compartments (Figure 1C).^34^ Megakaryocytic, erythroid, and lymphoid progenitors were delineated by expression of CD41, CD71, and CD127, respectively.^5,35^ Markers such as FcεRIa, CD170/Siglec-F, Ly-6G, and CD115/CD85k contributed to the classification of progenitors from the basophil-mast, eosinophil, neutrophil, and monocyte-dendritic lineage branches, respectively (Figure 1C- D).^36,37^

We aligned these preliminary cluster assignments with established transcriptional signatures for definitive annotation. Clusters annotated as HSCs and multipotent progenitors (MPPs) expressed high levels of *Hlf*/*Gcnt2*/*Meis1,* while megakaryocyte progenitors (MkPs) and megakaryocyte-erythroid progenitors (MEPs) showed enrichment of *Pf4*/*Vwf*/*Rab27b* and *Klf1/Ermap/Car1*, respectively (Figure 1E).^5,26,35,38–42^ Common myeloid progenitors (CMPs) and granulocyte macrophage progenitors (GMPs) shared classical myeloid transcripts *Mpo* and *Mif,* whereas the neutrophil lineage showed elevated amounts of *Ms4a3/Trem3/Ngp* within myeloid lineages (Figure 1E).^27,28,38^ Finally, high expression of *S100a4*/*S100a6* was detectable in the myeloid-DC compartment, whereas *Tcf4*/*Runx2* marked pDC-lineage committed progenitors (Figure 1E).

Comprehensive examination of the BM progenitor populations revealed a continuum of transcriptional states bridged by trajectory branches (Figure 1F). Pseudotime inference corroborated the Monocle3 predicted trajectories. We observed a closed-loop trajectory in the oligopotent progenitor compartment (CMP, GMP, MDP, cMoP (common monocyte progenitor), and CDP), supporting the concept of a highly plastic hematopoietic tree (Figure 1F). This pseudotime analysis highlighted a gap between clusters for several lineage branches, e.g. pre- Neutro to Neutro-I (Figure 1F), consistent with extensive transcriptional volatility during the transition of progenitors to mature cell states.

For interrogating surface markers, genes, TFs, and transcript-protein expression coherence of this novel dataset at single-cell resolution, the interactive online tool Progeniverse was developed and is freely available (https://sch197.med.uni-rostock.de/CITE-seq/mouse/BM-progenitors/).

### Immunophenotypic heterogeneity in BM progenitor compartments defined by a multimodal approach

Traditionally, identification of HSCs and progenitor cells relied on antibody panels, though overlapping surface marker patterns limit unambiguous sorting for subsequent functional analyses.^43^ Standard markers e.g., CD34 and FcγR, fail to distinguish progenitors of megakaryocyte, erythrocyte, DC, and basophil lineages, often grouped together as CMPs, masking actual population heterogeneity.^5,39^ Therefore, we applied unbiased analytical phenotyping to define potential novel marker combinations for optimized identification of hematopoiesis at single-cell level.

To identify decisive cell surface markers for CITE-seq immunophenotyping of HSPCs, we applied two main criteria: the marker (1) showed expression above background levels, validated by isotype controls, and (2) distinguished at least two HSPC states or lineages. In total, 97 markers fulfilled these conditions (Figure 2A; supplemental Figure 2 and 3).

**Figure 2.**
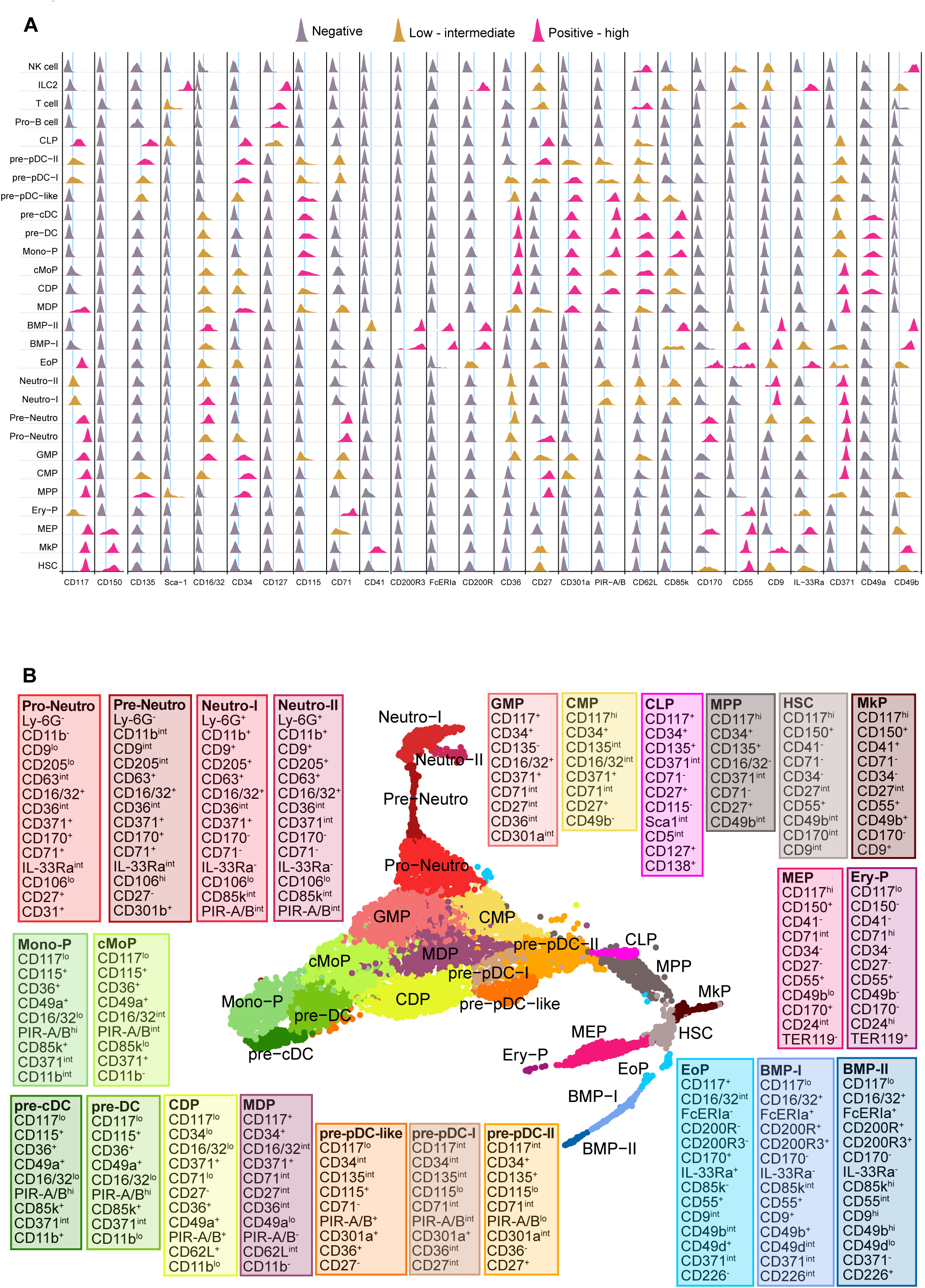
An optimized surface marker panel for detailed bone marrow progenitor profiling. (A) Ridge-plot depicting the surface level expression of 26 HSPC differentiation markers. These markers are nominated based on deferential abundance in one or multiple progenitor stages. Expression value of each ADT in each single cell is scale transformed to 0-1 (min-max) to calculate the ridge. Vertical black lines are used to separate the surface markers from each other and colored lines indicate the background threshold for each marker. Level of expression is indicated with color-codes as follows: gray = negatives; Orange = low-intermediate; magenta = positive-high. (B) Differentially enriched surface markers across lineages and transcriptionally closer progenitor stages suggested as cell-state-specific surface marker panel in color-matched boxes.

**Figure 3.**
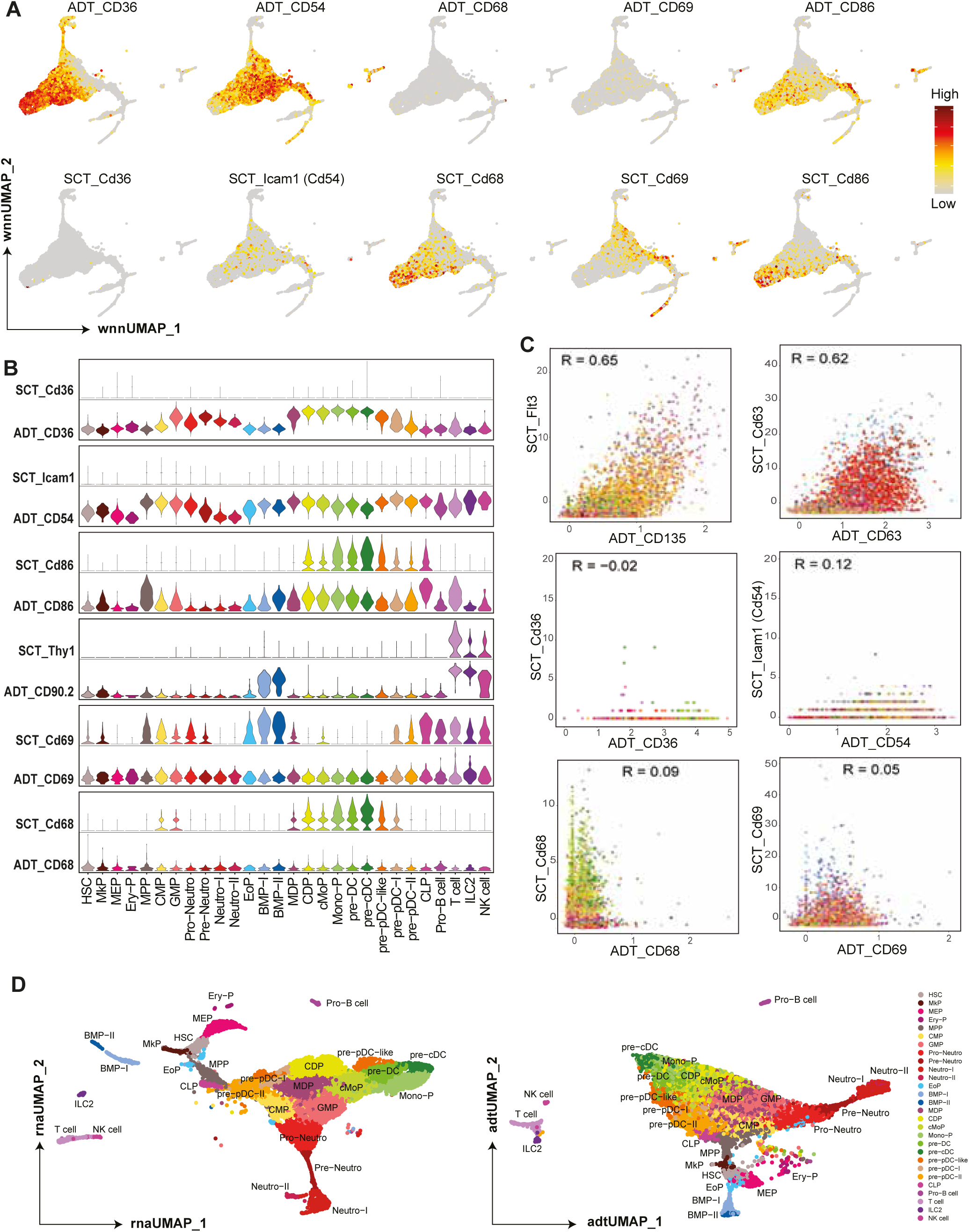
Discrepancies between surface protein versus transcript level expression in BM progenitor populations. (A) wnnUMAP projected normalized expression of five widely used hematopoietic cellular state identifying markers at ADT-measured surface protein level (top) and at corresponding transcript level (bottom). (B) Violin plots comparing the transcript level expressions with the corresponding ADT- measured surface protein expressions across the progenitor stages. Each surface protein is grouped together with the corresponding transcript. (C) FeatureScatter plots depicting the direct ADT versus transcript level comparisons of the respective hematopoietic cell surface markers. Pearson correlation coefficient values ‘R’ indicate the transcription-translation consistency of each marker. (D) Cluster neighborhoods visualized based on single modalities: transcriptional similarity in rnaUMAP (left) and surface protein similarity in adtUMAP (right). Cells are colored by cluster.

The combination of sample depth, progenitor enrichment, and a comprehensive surface marker panel enabled us to define novel specific combinations of surface proteins for almost all individual progenitor stages. For example, in the lin^-^ compartment, early progenitor clusters such as HSCs, MkPs and MEPs were identified as CD117^hi^CD150^+^CD34^-^, with CD41 and CD71 expression defining MkPs and MEPs, respectively, while HSCs lacked both. During the transition of the MEP towards the Ery-P stage, CD71/Ter119 levels progressively increased while CD117/CD150/CD170 expression declined. The consensus marker profile grouped MPPs, CMPs, and GMPs together as CD117^hi^CD150^-^CD34^+^ while relative expression of CD135 and CD16/32 further segregated this group. Common lymphocyte progenitors (CLPs), which are commonly defined as Lin^-^CD117^int^CD34^+^CD135^+^CD127^+^, were also enriched for CD138/CD27 expression (Figure 2A; supplemental Figure 2).

Expression of FcεRIa/CD200R/CD200R3 delineated the basophil-mast cell lineage stages BMP-I and BMP-II, which additionally exhibited high levels of CD85k/CD49b/CD90.2/CD226. Myeloid surface markers CD63 and CD301a captured two large subsets, neutrophils and monocyte-DC lineage progenitors. The subset of monocyte-DC progenitors, categorized as CD117^lo^CD115^+^CD301a^+^, also expressed CD49a/CD62L/CD85k. CD11b expression was first detectable within CDPs and was stepwise upregulated throughout successive developmental stages (Figure 2A; supplemental Figure 2). Based on the surface marker patterns observed here, we present an optimized immunophenotyping panel for high-resolution BM progenitor profiling (Figure 2B). Additionally, the full spectrum of CD surface-antigens was screened to identify non-ADT lineage-associated markers enriched at transcript level across progenitor stages (supplemental Figure 4).

In hematopoietic cells, transcript abundance poorly predicts the surface protein expression level, with only about 40% of the variance in protein levels is explained by mRNA levels, as the latter are influenced by several factors like post-translational modifications, protein trafficking, stability, and proteolytic processing, as well as technical noise of the experiment_._^16,17,44,45^

When assessing the relationship between mRNA and protein for widely used immunophenotypic markers by analyzing the ADT-to-gene correlation, several discordances were observed. For example, CD36 and CD270 showed robust protein expression, though they were below transcript detection levels across all lineages (Figure 3A; supplemental Figure 5). In contrast, CD68 and CD69 were detectable at transcript level but not at surface protein level (Figure 3A). Lineage-specific differential expression correlation was also observed, e.g. transcript levels of CD90.1 (Thy1) corresponded with surface protein levels in lymphoid clusters but not in BMPs (Figure 3B). Consistent markers showed a Pearson correlation coefficient of 0.50 or higher (e.g., CD135, CD63), whereas less consistent markers (e.g. CD68) showed 0.1-0.0 or even negative correlation values (CD36) (Figure 3C).

While lineage relationships were mostly conserved when comparing unimodal UMAPs based on transcripts (rnaUMAP) or ADTs (adtUMAP), subtle differences were observed, such as the pre-pDC-like population being separated into two subclusters in rnaUMAP, indicating their transcriptional lineage divergence not yet translated into surface profiles (Figure 3D).

These results caution against inferring functional states from surface markers or transcriptional profiles alone. Our comprehensive ADT-based CITE-seq immunophenotypic panel enables the redefinition of surface markers and suggests gating strategies for isolation and functional characterization of closely related progenitor stages.

### Mapping hematopoietic trajectories from the single-cell TF expression landscape

Single-cell analyses highlight a marked heterogeneity in TF expression across immune lineages, underscoring their role as key drivers in cell fate decisions, lineage transitions, and reprogramming potential.^14,19,46^

Integrating our dataset with information available from the mouse TF database AnimalTFDB 2.0^47^ identified differentially expressed TFs across progenitor populations, with patterns aligning to established lineage relationships. *Ebf1*, *Tcf7*, *Gata3,* and *Tbx21* specifically marked Pro-B cells, T cells, ILC2s, and NK cells, respectively (Figure 4A).^35^ *Hivep2*/*Ikzf3* were co-expressed across the lymphoid compartment. Consistent with their surface marker profiles, MkPs and MEPs also shared HSC-associated TFs (*Tal1*/*Gata1*/*Camta1*), while *Gfi1b* expression indicated early megakaryocytic/erythroid commitment of the HSCs.^35^ MkPs (LKCD150^+^CD41^+^) were highly enriched for *Prdm5*, *Vezf1*, and *Pbx3* (Figure 4A).

**Figure 4.**
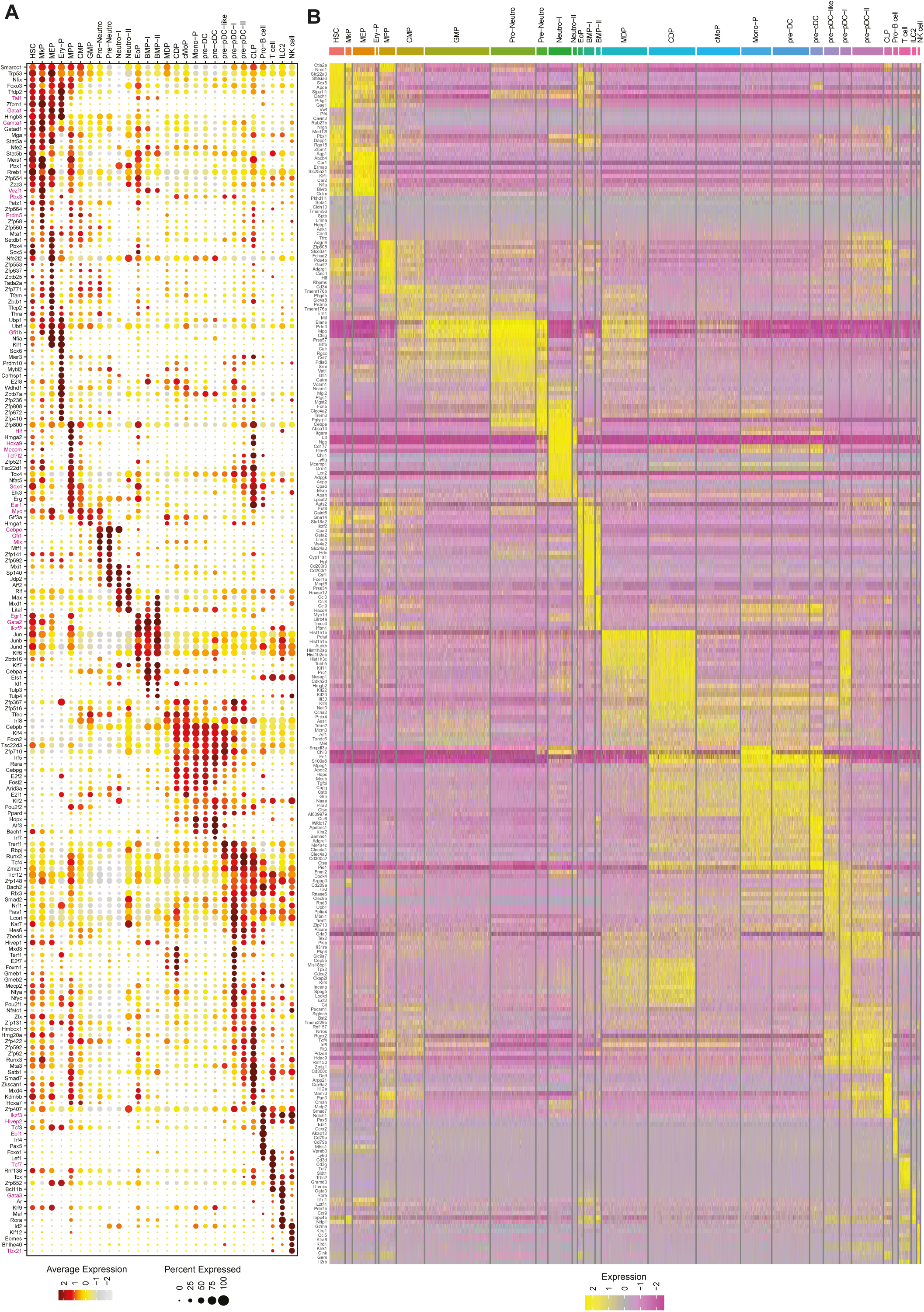
Cluster-specific gene enrichment analysis defines a comprehensive BM progenitor stage-specific transcriptional signature. (A) Bead plot showing the expression of signature TFs in BM hematopoietic progenitor stages. Mean expression is represented as a color scale of relative intensities, and the size of the circle represents the fraction of cells within that cluster having non-zero expression values. Example TFs mentioned in the result section are color-coded in magenta. (B) Expression of selected cluster-specific signature genes in the annotated hematopoietic cell states. 272 cluster-specific enriched genes were nominated from 28 clusters and mapped across 9281 single cells.

The multipotent progenitor compartment displayed a core set of TFs characteristic of uncommitted progenitors, such as *Myc*, *Esr1*, and *Sox4,* which were present in MPPs, CMPs, GMPs, and CLPs without indicating lineage commitment. Early hematopoietic regulators *Hlf* and *Mecom* supported annotation of MPPs, while shared expression of *Hoxa9* and *Tcf7l2* positioned them at an ancestral stage of CLPs.^48^ Neutrophil-defining TFs, including *Cebpe*, *Gfi1*, and *Mlx,* emerged during GMP-to-Pro-neutrophil transition but were absent from HSCs- to-early myeloid stages. In contrast, TFs defining eosinophil, basophil and mast cell progenitors, e.g. *Gata2*, *Ikzf2*, and *Egr1,* appeared in early HSCs and MPPs but not in CMPs or GMPs (Figure 4A).

We selected 272 cluster-specific signature genes and visualized their transcript profiles across 28 clusters in a heatmap (Figure 4B). This revealed a strong transcriptional relationship of MDPs with CDPs, though they are phenotypically closest to the GMPs (Figure 2A; Figure 4B).

This network of hematopoietic differentiation and lineage-determining key TF reveals dynamic regulatory relationships.

### CITE-seq reveals five distinct neutrophil committed progenitor clusters in mouse BM

Neutrophils are the most abundant peripheral blood leukocytes and require continuous replenishment from BM due to their short lifespan.^49–51^ Despite known differences in nuclear morphology, granule composition, and functional capacity across maturation stages, their developmental landscape remains incompletely defined, partly due to their low transcript counts.^50,52^ Using CITE-seq, we identified 5 novel and distinct developmental stages from GMP to mature neutrophils, each defined by unique surface markers and TFs. Among these, early granulocytic progenitors, GMPs and Pro-neutrophils represent a significant portion of the hematopoietic progenitor pool (Figure 5A).

**Figure 5.**
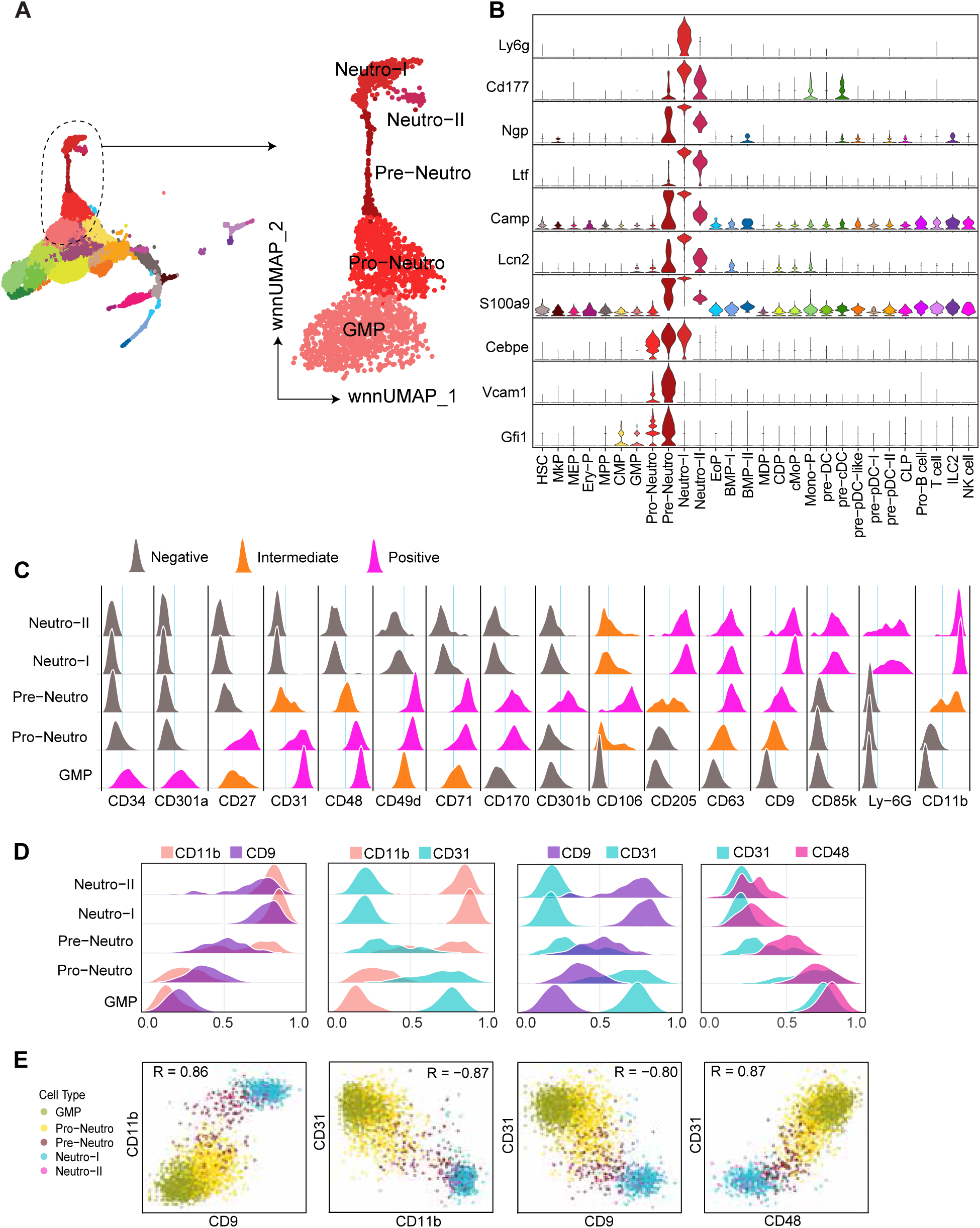
Multimodal marker identification dissects transitional neutrophil lineage stages. (A) wnnUMAP showing the neutrophil lineage populations starting from GMP stage. (B) Violin plots depicting neutrophil lineage specific TFs, signature transcripts, and immune effector genes. (C) Differentially expressed cell surface markers across neutrophil lineage stages. Level of expression is indicated with color-codes as follows: gray = negatives; Orange = intermediate; magenta = positive. (D) Overlaid ridge-plots illustrating the co-expression pattern of hematopoietic progenitor marker CD9, CD31, and CD48 with the neutrophil maturation marker CD11b. Cell surface markers are color-coded according to the respective legends on the top of each plot. (E) FeatureScatter plots showing the distribution of neutrophil lineage stages with different combinations of neutrophil cell surface markers CD11b, CD9, CD31, and CD48.

TF repertoire showed that both mature and progenitor neutrophil clusters expressed *Cebpe,* consistent with their lineage identity. Immature neutrophil progenitors were enriched for the TF *Gfi1,* which represses alternative lineages and promotes neutrophil maturation. *S100a9* was enriched starting from the pre-Neutrophil stage, suggesting a possible role in neutrophil maturation (Figure 5B). More mature neutrophil populations showed enrichment for secreted proteins involved in immune activity, such as *Ltf, Ngp, Lcn2*, or *Cd177* (Figure 5B). Unsupervised wnnUMAP revealed two mature neutrophil populations sharing similar surface markers despite striking transcriptional divergence. The expression level of the hallmark neutrophil maturation marker Ly-6G was comparable in both stages, though *Ly6g* transcription was already curtailed in Neutro-II. Reduced expression of immune effector genes, such as *Camp* or *Cxcr4* implies that Neutro-II cells may constitute a marginated pool for emergency release (Figure 5B-C).^49^

Having examined the transcriptional profile, we sought to identify specific surface marker patterns of the neutrophil progenitor populations. We observed a gradual repression of progenitor associated markers (CD34/CD48/CD31/CD49d) and escalation of neutrophil lineage-associated markers (CD11b/Ly-6G/CD9/CD63/CD205) across the neutrophil progenitor stages. Strikingly, pre-Neutrophils displayed peak levels of CD106 on the surface and its corresponding transcript, *Vcam1*, underscoring its role in promoting adhesion to BM stromal cells and endothelium, a prerequisite for proper retention and timely release of mature neutrophils into the circulation (Figure 5C). Notably, CD71 and CD170 showed transitional upregulation in pro- and pre-neutrophils but declined as the cells matured (Figure 5C). Progenitor markers CD9/CD71 outperform traditionally used neutrophil-defining markers CD11b/CD63 to track the lineage commitment and maturation, as the latter two showed broader expression (Figure 5D-E; supplemental Figure 2).

Despite diverse biological functions, our data defines a single continuum of neutrophil development in the mouse BM.

### pDC progenitor populations share a complex lineage affiliation

pDCs are immune sentinels producing type I interferons in response to viral nucleic acids and, to a lesser extent, presenting antigens.^53,54^ Their developmental ontogeny remains debated, with evidence supporting both myeloid and lymphoid origins.^55–59^ Recent high-dimensional studies reported an unexpected pDC heterogeneity^60^, a pDC-to-cDC fate-switching mechanism^61^, and a novel transitional DC (tDC) population with a yet undefined ontogeny^62^.

In CITE-Seq profiling, all previously transcriptionally defined pDC progenitor clusters (pre-pDC-like, pre-pDC-I, and pre-pDC-II) exhibited a unique transcriptional and surface marker profile, setting them apart from monocyte-DC and lymphoid progenitors and supporting their classification as a distinct lineage branch (Figure 6A). pDC progenitors differed from classical monocyte-DC progenitors by exhibiting higher levels of early hematopoietic markers CD117/CD34, coupled with diminished expression of established monocyte-DC markers (CD115/CD85k), indicative of an early lineage divergence (Figure 6B). Despite shared expression of markers like CD301a and CD371, other typical monocyte-DC markers, including CD49a and CD62L, were reduced or absent in pDC progenitors (Figures 2A and 6B). Phenotypic heterogeneity within the pDC progenitor continuum was evident, with a gradual upregulation of the lymphoid-associated marker CD27 and downregulation of myeloid markers CD36 and PIR-A/B from pre-pDC-like to pre-pDC-II, implying either a dual lympho-myeloid origin or bifurcating trajectories toward pDCs and cDCs (Figure 6B). Comparing CD27 against CD36 or PIR-A/B could be a useful strategy to phenotypically define these pDC progenitor stages for functional implications (Figure 6C-D).

**Figure 6.**
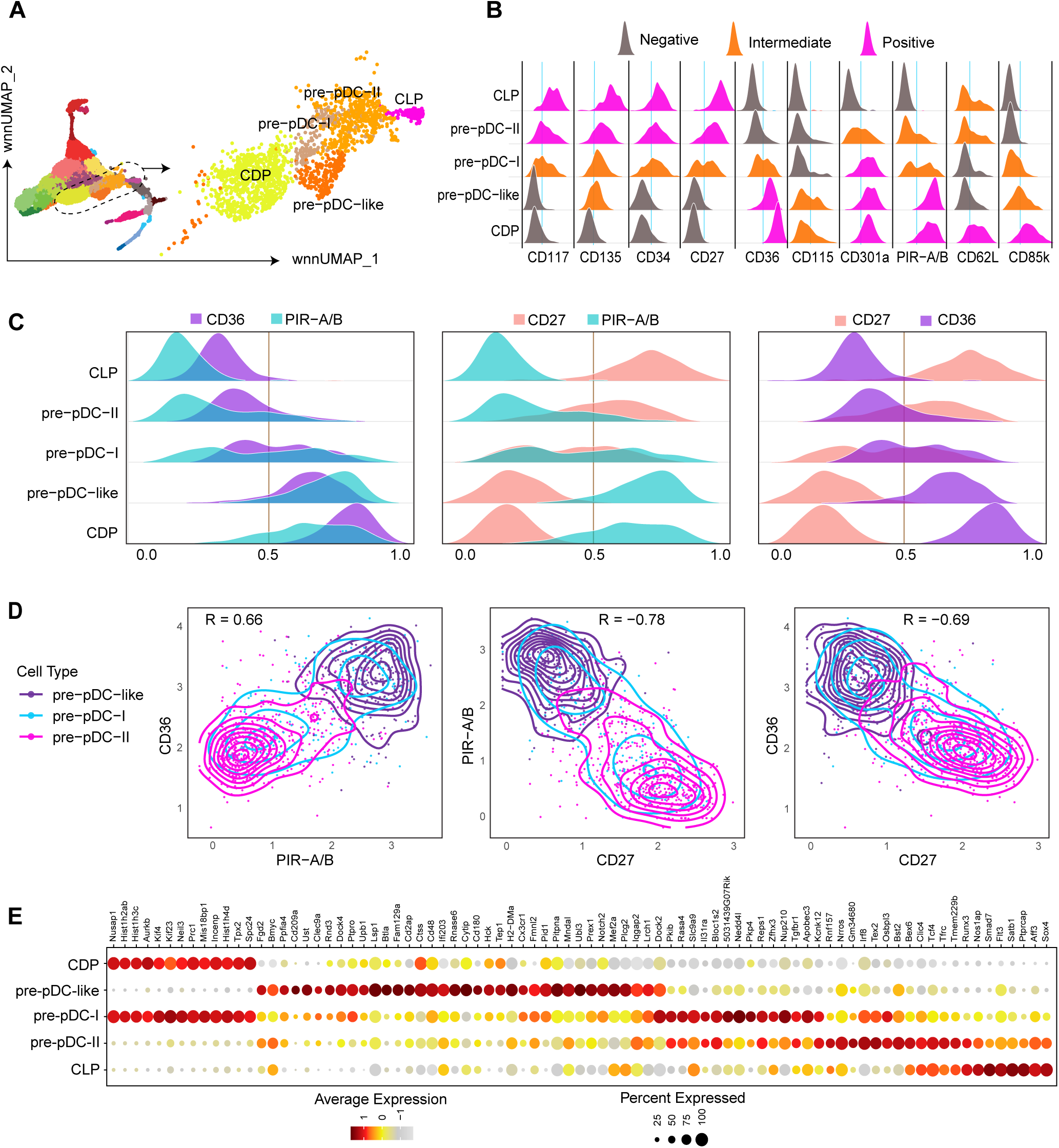
pDC progenitors follow a complex hematopoietic lineage tract. (A) wnnUMAP showing the pDC progenitor stages together with common DC progenitors (CDPs) and common lymphoid progenitors (CLPs). (B) Surface marker expression pattern across the pDC progenitor clusters compared to CDPs and CLPs. Level of expression is indicated with color-codes as follows: gray = negatives; Orange = intermediate; magenta = positive. (C) Overlaid ridge-plots illustrating co-expression pattern of surface marker CD36, PIR-A/B, and CD27 across pDC progenitor stages. (D) FeatureScatter plots showing the distribution of pDC progenitor populations according to the ADT-derived expression levels of these cell surface markers CD36, PIR-A/B, and CD27. (E) Bead plot of transcriptional signatures across pDC progenitor clusters as well as CDPs and CLPs. Here the size of the circle is the fraction of cells and the color represents mean expression level.

Transcriptional signatures followed the same trend. Both pre-pDC-I & II clusters showed strong association with mature pDC hallmark transcripts *Bst2*, *Flt3*, *Irf8*, and *Tex2*. CD27^+^CD36^lo^ pre- pDC-II showed expression of CLP markers such as *Nos1ap*, *Sox4*, and *Satb1,* suggesting a probable lymphoid origin of these cells from CLPs. In contrast, CD27^int^CD36^int^ pre-pDC-I showed extensive transcriptional overlap with CDPs such as *Tpx2*, *Spc24*, and *Neil3,* establishing them as myeloid-derived pDC progenitors (Figure 6E). On the other hand, weaker expression of pDC hallmark transcripts positioned the pre-pDC-like cells at an earlier developmental stage than that of pre-pDCs. Top expressed genes in CD27^-^CD36^+^ pre-pDC-like cluster included factors of both pDC and cDC specification, such as *Notch2*/*Ubl3*/*Mef2a.* Thus, though these cells are biased towards the DC lineage, they still possess a dual potential and may represent tDC progenitors (Figure 6E).^62,63^ Of note, at least 50% of pre-pDC-like cells showed high expression of DC-specific receptors such as *Clec9a*^64^ and *Cd209a* (DC-SIGN), indicating a presence of cDC1 progenitors here (Figure 6E).

Our analysis revealed the transcriptional and surface marker heterogeneity among the pDC progenitor stages with a broad overlap of features of pre-pDC-II with CLPs and pre-pDC-I with CDPs, where the pre-pDC-like cluster represented bipotent features.

### A CITE-seq derived gating approach for rare hematopoietic progenitor populations

Isolating transcriptome-defined hematopoietic progenitor stages as pure populations is a prerequisite for future function- and fate-mapping. A hematopoietic binary logic tree showed that 16 prospective progenitor markers could effectively lead via binary decisions to 21 progenitor stages (Figure 7A). We projected these progenitor populations in bi-axial ADT-defined scatterplots and calculated the distribution of the respective populations (Figure 7B). Using a 20-surface marker panel, we schematically illustrated the stepwise isolation of 21 distinct progenitor populations as a flow cytometry-like gating strategy. Our analysis revealed that CD9/CD27/CD36/CD55/CD85k/CD106/PIR-A/B harbor a strong potential for defining mouse hematopoietic progenitor stages. We observed that CD55 marks eosinophil, basophil, and mast cell progenitors, whereas combined CD71/CD170 expression identified two transitional neutrophil progenitor stages, Pro- and Pre-neutrophils, which can be further separated using CD27 and CD106 (Figure 7B). As the expression pattern of individual surface markers may vary under different biological conditions, we proposed lineage- and stage-specific alternative markers. Specifically, CD127 could be substituted with CD138 to distinguish CLPs, PIR-A/B with CD36 to define pDC lineage progenitors, and CD106 with CD31 within the neutrophil lineage (supplemental Figure 6).

**Figure 7.**
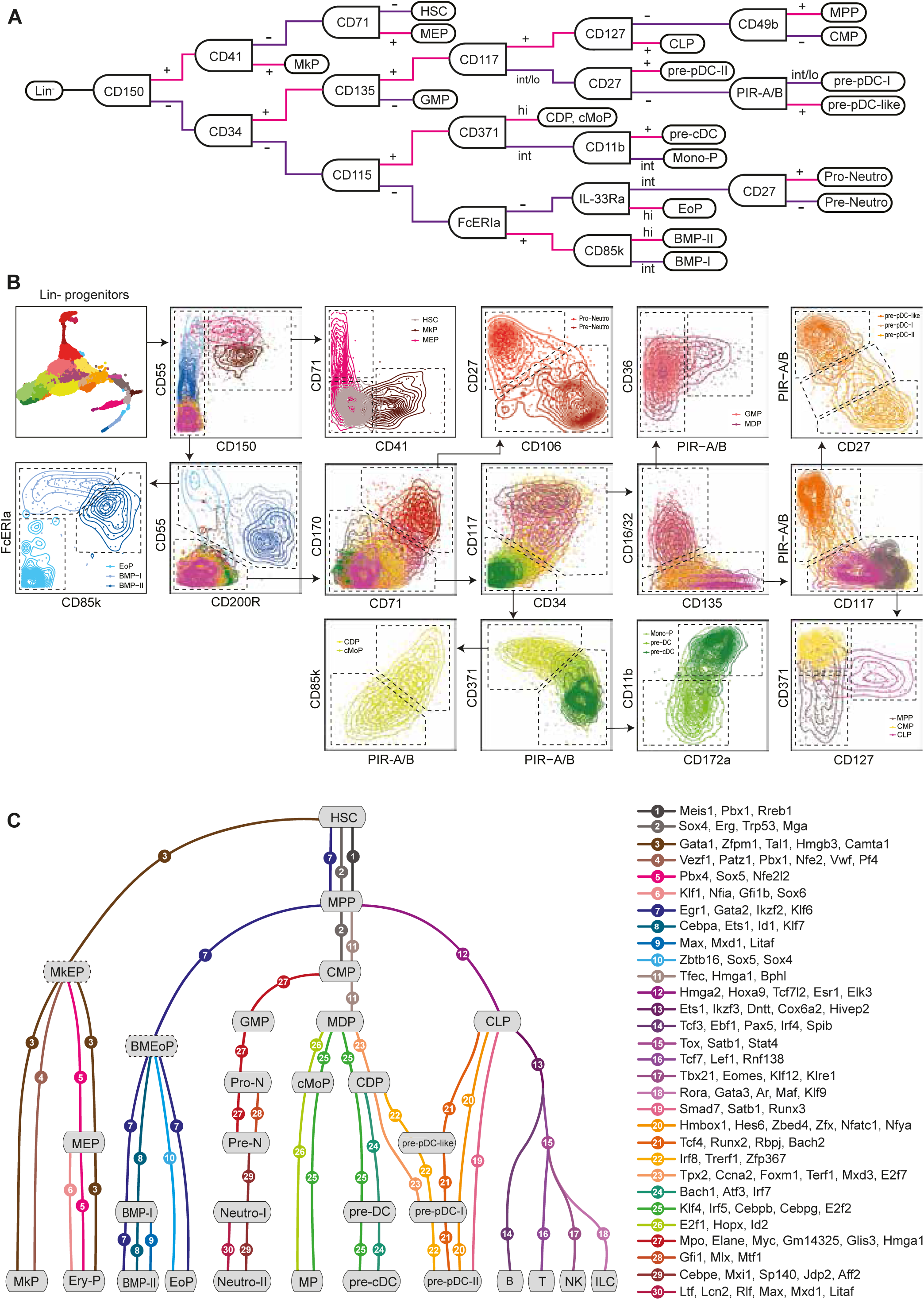
A refined classification of the mouse BM hematopoietic progenitors compartment identified novel surface markers and TFs network. (A) Hematopoietic binary logic tree showing cell surface markers representing prospective binary decision factors of lineage and stage segregation in the BM progenitors compartment. (B) A *posteriori* sequential gating strategy for the definition of 21 CITE-seq derived BM lin^-^ progenitor stages to emulate flow cytometry analysis. A 20-markers panel is used to show the schematic isolation strategy. Dotted lines are used to show the proposed gates. Respective cluster annotations are mentioned in terminal gates. Lin = B220, CD3, CD4, CD8a, CD11b, CD11c, CD19, Ly-6G, MHC-II, NK1.1, and Ter119. (C) A “subway map” representation of the TF regulatory landscape is constructed by wiring the lineages and progenitor stages together with the TF modules. 28 identified and 2 proposed (dotted border) progenitor stages are represented as subway stations and 30 TF modules are used as driving factors to connect two or more hematopoietic stages as subway lines.

Finally, to facilitate the integration of other transcriptomic datasets with our proposed surface marker panel and gating scheme, we constructed a comprehensive hematopoietic TF-network by wiring the lineages and progenitor stages with TF-modules composed of 3-6 TFs co-expressed at elevated levels. We projected the TF modules in a “subway map” comprising thirty TF modules driving the differentiation/developmental choices of 28 cellular stages from eight different lineages (Figure 7C). As expected, erythroid and megakaryocytes share a large spectrum of TFs (module-3: Gata1-Zfpm1-Tal1-Hmgb3-Camta1) throughout the whole developmental stages from HSCs to MkPs and Ery-Ps, though the stage defined as MEP only share TFs with Ery-Ps. We thus propose the existence of an additional common stage in between that we termed as MkEP. We made similar observations for Basophil-Mast cell and Eosinophil lineages as well, where module-7 (Egr1-Gata2-Ikzf2-Klf6) drives their development through HSC and MPP stages, though we did not observe any common stage (proposed as BMEoP) downstream to MPP (Figure 7C). TF kinetics of the pDC progenitor clusters represent their complex lympho-myeloid lineage affiliation, where transcriptional specification started latest from CLP and MDP stages, respectively.

Our data-driven gating approach, together with the identified TF-module network might permit effective isolation of transcriptionally defined mouse hematopoietic progenitor stages for more precise functional elucidations.

## 5. Discussion

Detailed characterization and isolation of hematopoietic stem and progenitor states hold the promise for advancing therapies across a range of hematologic and non-hematologic disorders. Recent findings from several studies have challenged the precise identification of hematopoietic progenitor compartments based on single-cell transcriptomes or flow-cytometry, alone. While flow-cytometry provides an incomplete biological landscape, single-cell transcriptomic profiling is not fully applicable in clinical diagnostics. The combined application of complementary high-throughput multimodal analyses of the BM progenitor compartment generated a detailed map connecting dynamic transcriptional states with corresponding cell surface markers.

Here, we report a 123-ADT CITE-seq panel integrated with single-cell TFs profiling of 9,281 rare mouse BM HSPCs to facilitate the prospective identification and fate mapping of distinct hematopoietic progenitor stages. In contrast to scRNA-seq, ADT counts faithfully represent surface marker abundance, thereby permitting a quantitative alignment with flow cytometry. Most of the conventional progenitor markers, including CD117, CD34, CD115, and CD127, mapped closely to transcriptionally defined clusters. However, other markers such as CD43, CD44, CD48, and Ly-6C showed overlapping expression among lineages and demand additional attention for future cytometry-based applications. We annotated the mono-DC progenitors as CD117^lo^, while traditionally flow cytometry describes them as CD117^int/lo^. Note that CD117 was spiked into the CITE-seq cocktail with a pre-titrated dilution of 1:1600 due to its high expression level among BM progenitors. On the other hand, basal high ADT detection was observed for CD135 (Flt3), which was spiked-in with 1:100 dilution due to its low expression pattern in flow cytometry-based pre-titration. Despite high basal expression, the CD135 ADT profile matched the expected phenotype of MPPs, CMPs, CLPs, as well as pDC progenitors, but was indistinguishable in the classical DC progenitor stages. Notably, inconsistencies between surface and transcript expression levels, as observed for CD36, CD54, and CD69, indicate scRNA-seq might often fail to detect low-abundance transcripts due to technical dropouts, though the corresponding protein may be expressed. It also highlights that scRNA-seq cannot reflect possible mRNA degradation, alternative splicing, microRNA repression, or retained protein localization and trafficking.

Our inclusive approach to sort lin^-^ cells enabled us to capture a comprehensive spectrum of stem and progenitor compartments comprising eight hematopoietic lineages leading to refined insights into trajectories and branching sites. In contrast to existing models, the progenitor stage we captured as MEPs lacked megakaryocytic markers and exclusively expressed erythroid markers (CD71/CD55) instead, suggesting it might be more accurately classified as early erythrocyte progenitors or pre-Ery-P, a conclusion also supported by other recent studies.^39^ Of note, we could not rule out a possible shared stage for these two lineages (MkEP) owing to the fact that they share many TFs. Within the granulocyte compartment, neutrophil progenitors exhibited a stepwise restricted TF profile from the GMP to mature stages, whereas the eosinophil, basophil, and mast cell progenitors showed TF profiles shared with multipotent stages, indicating an early lineage segregation from MPPs without transiting through CMP or GMP intermediates.

Importantly, our study establishes a framework for functional profiling of the hematopoietic progenitor compartments beyond the identification of certain states. At our progenitor-enriched resolution, we identified two discrete stages between GMPs and mature neutrophils, both are highly distinct in their surface marker and TF profiles. Neutrophil-II with reduced immune effector expression may represent a marginated or quiescent back-up neutrophil pool, which may further contribute to understanding the rapid turnover during granulopoiesis. Whereas the ontogeny and developmental relationship of pDC progenitors have been debated for decades, our dataset reveals that, in addition to the core pDC signatures, early pDC progenitors are heterogeneous in both TF and surface marker profiles, exhibiting a great overlap with lymphoid and mono-DC lineage stages. This highlights their possible lympho-myeloid shared origin and/or putative bipotent fate in health and disease. We report that pDC progenitor populations do not fit perfectly within either of these two lineages. Instead, we suggest that together with the potential tDC progenitors, pDC progenitor populations would be best classified as a separate lineage branch.

This work introduces a digital atlas of BM progenitors, featuring an RNA versus ADT viewer as a specifically tailored online tool, to support the community in exploring their own data, comparing the molecular regulators, and refining progenitor identification strategies. Wider adoption of this strategy has the potential to minimize conflicting outcomes due to imprecise population definitions and thus to reconstruct the hematopoietic hierarchies by resolving the underlying regulatory networks involved in progenitor cell fate choices.

Last, this extensively annotated and curated dataset provides a valuable reference resource for future discoveries in both hematology and immunology. We anticipate that this integration of surface epitope expression together with single-cell transcriptomes and the proposed novel gating scheme and molecular TF wiring of the stages developed from that, will clarify many unresolved aspects of BM progenitor cell states under diverse biological conditions.

## 6. Acknowledgments

The authors thank Sonja Schavier at the Institute of Medical Microbiology and Hospital Hygiene, Medical Faculty and University Hospital Düsseldorf for her help with the mouse bone marrow preparation. Stefanie Scheu was supported by the Deutsche Forschungsgemeinschaft (DFG, German Research Foundation) projects SCHE692/6–1, SCHE692/8–1, and DFG- 270650915/GRK2158 and the Manchot Graduate School ‘Molecules of Infection IV’.

## 7. Authorship Contributions

M. H. B., S. A., and S. S. conceptualized the study; M. H. B., and S. A. conducted experiments; T. L. performed the sequencing; T.L., and S. B. curated the data; M. H. B., S. B., and F. S. visualized the data; F. S., M. B. H., and S. S. constructed the data visualization portal; M. H. B., S. B., N.G., and S. S. interpreted data, wrote the original draft, reviewed and edited the manuscript; and S. S. supervised the study; all authors reviewed and approved the manuscript.

## 8. Conflict of Interest Disclosures

S. B. is presently employed by Novartis, Basel, Switzerland, which is unrelated to this work. The remaining authors declare no competing financial interest.

### Data sharing statement

The data reported in this study have been deposited in the European Nucleotide Archive (ENA) database under accession number PRJEB109753 (https://www.ebi.ac.uk/ena/browser/view/PRJEB109753). The processed single-cell CITE-seq data have been deposited in Zenodo (https://doi.org/10.5281/zenodo.19696575). All codes used in this study are available in Github repository: https://github.com/Mahamudul-Bhuyan/CITE-seq_Mouse_BM_progenitors. This BM HSPC data were also uploaded to an online visualization platform: https://sch197.med.uni-rostock.de/CITE-seq/mouse/BM-progenitors/

## Visual Abstract

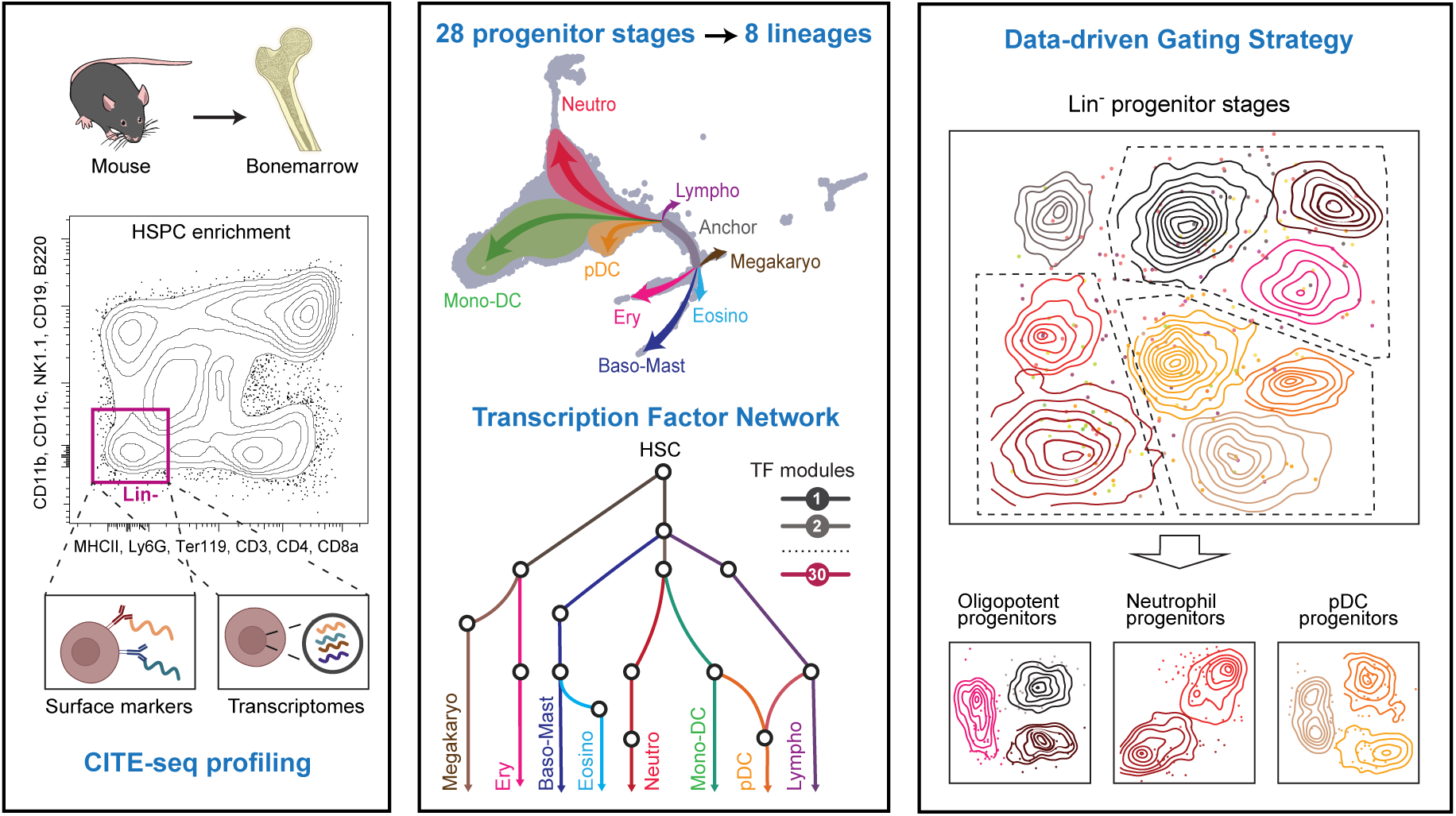

