## Supplemental Materials for "An immunophenotype-coupled transcriptomic atlas of mouse hematopoietic progenitors"


**Authors and affiliations:**

Mahamudul Hasan Bhuyan,[1,2] Sumanta Barman,[3] Shafaqat Ali,[1,2] Felix Steinbeck,[1] Tobias Lautwein,[4] Norbert Goebels,[3] Stefanie Scheu[1,2]

[1]Institute of Immunology, Rostock University Medical Center, Rostock, Germany

[2]Institute of Medical Microbiology and Hospital Hygiene, Medical Faculty and University Hospital Düsseldorf, Heinrich Heine University Düsseldorf, Germany

[3]Department of Neurology, Medical Faculty and University Hospital Düsseldorf, Heinrich Heine University Düsseldorf, Germany

[4]Biological and Medical Research Center (BMFZ), Heinrich Heine University Düsseldorf, Düsseldorf, Germany

**Corresponding author**

Prof. Dr. Stefanie Scheu

Institute of Immunology, Rostock University Medical Center

Schillingallee 70, 18057 Rostock, Germany


**Content**


Supplemental Methods

Supplemental Figures 1-6

Supplemental Tables 1-4

Supplemental References

### Supplemental Methods

#### Preparation of BM and cell sorting

Lin$^-$ HSPCs were enriched from the BM of six 10-weeks-old C57BL/6 mice. BM was extracted from the femurs and tibias via centrifugation method[1] and lineage depletion was done by FACS sorting to enrich lin$^-$ HSPCs using the negative selection against the surface markers CD3, CD4, CD8a, CD11b, CD11c, CD19, B220, MHC-II, Ly-6G, NK1.1, and Ter119 (supplemental Table 1). 7-amino-actinomycin-D (7-AAD) was used for dead cell exclusion. Cell sorting was performed using BD FACSAria III (BD Biosciences). Detailed gating strategy is shown in supplemental Figure 1.

#### Sample preparation for CITE-seq

Cell hashing was performed by using TotalSeq™-A anti-mouse Hashtag antibodies 1-6 (supplemental Table 2) to tag the cells from each individual animal. After 3x washing, cells were pooled together for CITE-seq staining. TotalSeq™-A Mouse Universal Cocktail, v1.0 (119 antibodies and 9 isotypes) was used for CITE-seq staining (supplemental Table 3). According to the manufacturer's protocol, cells were stained in 50µl of volume. Four additional CITE-seq antibodies against the surface markers CD117, CD34, CD135, and CD16/32 were pre-titrated and spiked-in according to the manufacturer's protocol to the rehydrated totalSeq-A cocktail (supplemental Table 4). Cells were stained with this spiked-in cocktail for 30 min at 4°C. Before 10X chip loading, Trypan Blue staining and counting on a hemacytometer confirmed the viability over 90%.

#### Single-cell CITE-seq library generation and sequencing

Single-cell capture, cDNA library preparation, and sequencing were performed by the Biomedical Research Center sequencing core facility at Heinrich Heine University Duesseldorf, Duesseldorf, Germany. A total of 20,000 cells were used as input for the single-cell droplet library generation on the 10X Chromium Controller system, utilizing the Chromium Single Cell 3'NextGEM Reagent Kit v3.1 with Feature Barcode technology for Cell Surface Protein (protocol version CG000317) according to the manufacturer's instructions. Sequencing was carried out on a NextSeq 2000 system (Illumina Inc., San Diego, USA) with a mean sequencing depth of 50,000 reads/cell for the Gene Expression library. The ADT library was sequenced to a depth of ~15,000 reads/cell, and the Cell Hashing library was sequenced to a depth of ~5,000 reads/cell.

#### Processing of 10X Genomics single cell data

Raw sequencing data was processed using the 10X Genomics CellRanger software (v7.0). Raw BCL-files were demultiplexed and processed to Fastq-files using the CellRanger *mkfastq* pipeline. Alignment of reads to the mm10 genome and UMI counting was performed via the CellRanger *count* pipieline to generate a gene-barcode matrix. The Cellranger *aggr* pipeline was used for aggregation and sequencing depth normalization.

**Bioinformatic analysis of single-cell CITE-seq data**

Filtered feature-barcode matrix files were analyzed with the R package 'Seurat'[2] (v5.2.1). Data reliability was ensured through initial quality control steps, including analysis and visualization of RNA features per cell, RNA counts per cell, and mitochondrial proportions for each cell. FeatureScatter visualizations examined relationships between nCount_RNA and nFeature_RNA, as well as mitochondrial percentage. Cells exhibiting abnormally low or high RNA feature counts (UMI < 1000 or UMI > 40000) and/or elevated mitochondrial proportions (>5%) as indicators of low-quality cells or doublets were excluded.

RNA assay normalization was performed using SCTransform, which identified variable features, scaled and centered the dataset. PCA was applied using the RunPCA function to reduce dimensionality. Batch effect correction was performed using the IntegrateLayers function with HarmonyIntegration to account for technical variation, producing an integrated ‘harmony’ reduction. A shared nearest neighbor graph was constructed using the FindNeighbors function, incorporating the first 20 dimensions of the harmony reduction. Dimensional reduction for visualization was performed using Uniform Manifold Approximation and Projection (UMAP) via the RunUMAP function using the first 20 harmony dimensions.

ADT assay normalization was performed using centered log-ratio (CLR) transformation, and the data were scaled and centered. Cells were multiplexed using Hashtag Oligonucleotides (HTOs) to distinguish cells from each individual animal and two different CITE-seq reactions. HTO barcode counts were obtained from multimodal analysis in Cell Ranger. HTO demultiplexing was performed using the 'HTODemux' function in the Seurat package (v5.2.1), and cells were classified as singlets (a cell confidently assigned to a single hashtag), doublets (multiplets assigned to more than one hashtag), or negatives (no confident HTO assignment) based on normalized HTO signals. This quality control step retained 12,128 singlets out of 20,000 events, filtering out doublets and negatives. PCA was applied using the RunPCA function on all ADT features to reduce dimensionality. Batch effect correction was performed using the FindIntegrationAnchors() function with CCA, followed by the IntegrateData() function to account for technical variation. A shared nearest neighbor graph was constructed using the FindNeighbors function incorporating the first 18 principal components. Dimensional reduction for visualization was performed using UMAP via the RunUMAP function with 18 dimensions.

The transcriptomic (RNA) and ADT assays were merged into an integrated multi-assay analysis. Weighted nearest neighbor (WNN)[3] analysis was performed using the

FindMultiModalNeighbors function, with RNA harmony reduction (dimensions 1:20) and ADT PCA reduction (dimensions 1:18). WNN-based UMAP (wnnUMAP) was computed using RunUMAP on the weighted nearest neighbor graph. Cell clusters were identified using the FindClusters function on the WNN graph with a resolution parameter of 1.5, following the Seurat CITE-seq reference guide. The final filtered dataset was composed of 9,281 cells and 28 clusters containing 18,941 genes in total with a median detection of 4,477 genes per cell.

To prioritize oligonucleotide-conjugated CITE-seq antibodies in their immunophenotypic value, each transcriptionally defined cell cluster was manually annotated to the most likely cell population based on the specifically enriched ADTs derived from their surface markers. Underperforming ADTs were not considered for the manual cluster assignment, where it was defined by an indistinguishable staining pattern across the clusters. Corresponding transcript signatures were also used for the validation of the manual annotation.

Trajectory analysis was performed in R (version 4.4.3) with a Trajectory analysis package, Monocle3[4] (version 1.4.27). Trajectory analysis was embedded in wnnUMAP from surface markers and transcriptomic analysis. A pseudotime value was assigned to the cells in each branch with the Monocle3 pseudotime algorithms, with the root node set at the cluster identified as HSCs.

### Supplemental Figures

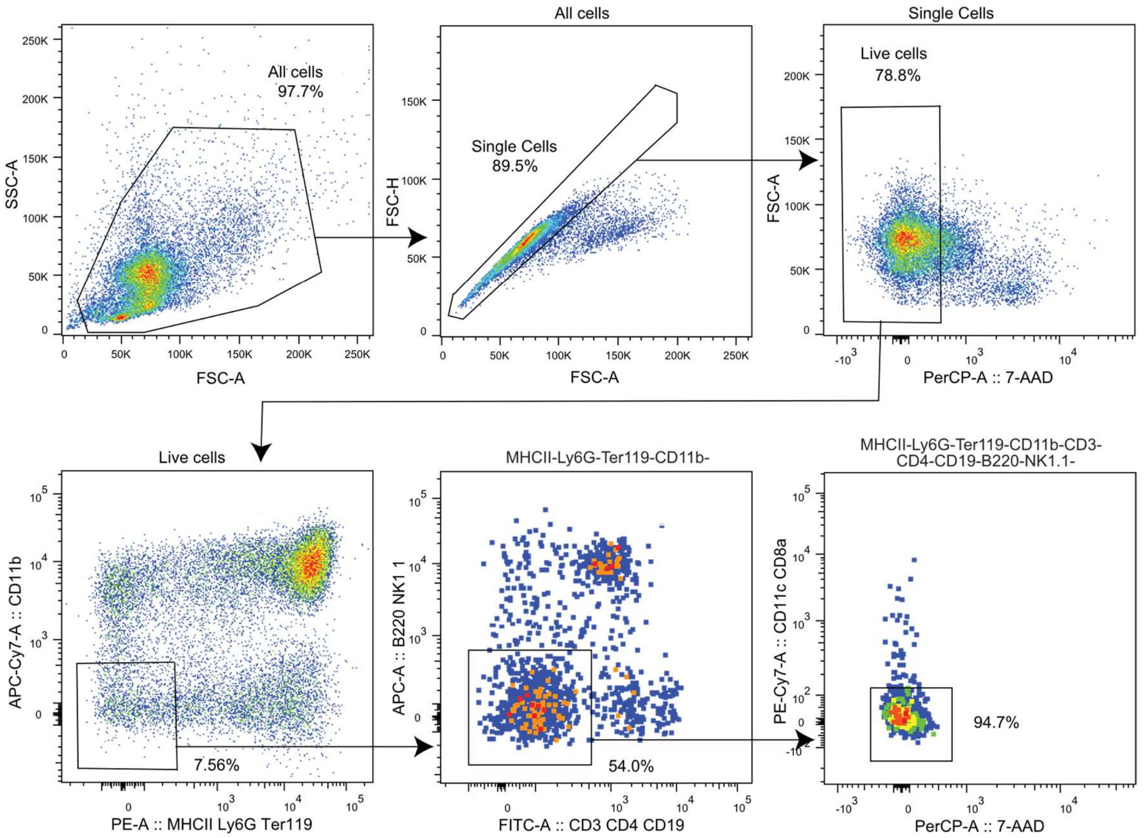


**Supplemental Figure 1:** Sorting strategy of BM lin$^{-}$ stem and progenitor cells. Viable cells were selected as 7-AAD$^{-}$ and sequentially gated negative for BM mature lineage markers: B220, CD3, CD4, CD8a, CD11b, CD11c, CD19, Ly-6G, MHC-II, NK1.1, and Ter119.

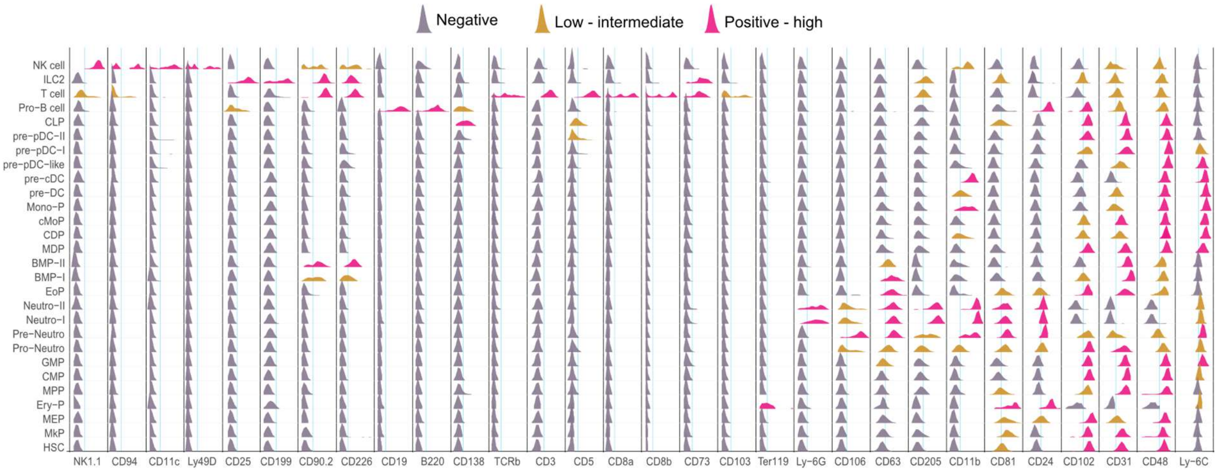


**Supplemental Figure 2:** Ridge-plot depicting the surface level expression of 30 mature lineage and progenitor markers. The expression value of each ADT in every single cell is scale transformed to 0-1 (min-max) to calculate the ridge. Vertical black lines are used to separate the surface markers from each other and colored lines indicate the background threshold for each marker. Level of expression is indicated with color-codes as follows: gray = negatives; Orange = low-intermediate; magenta = positive-high.

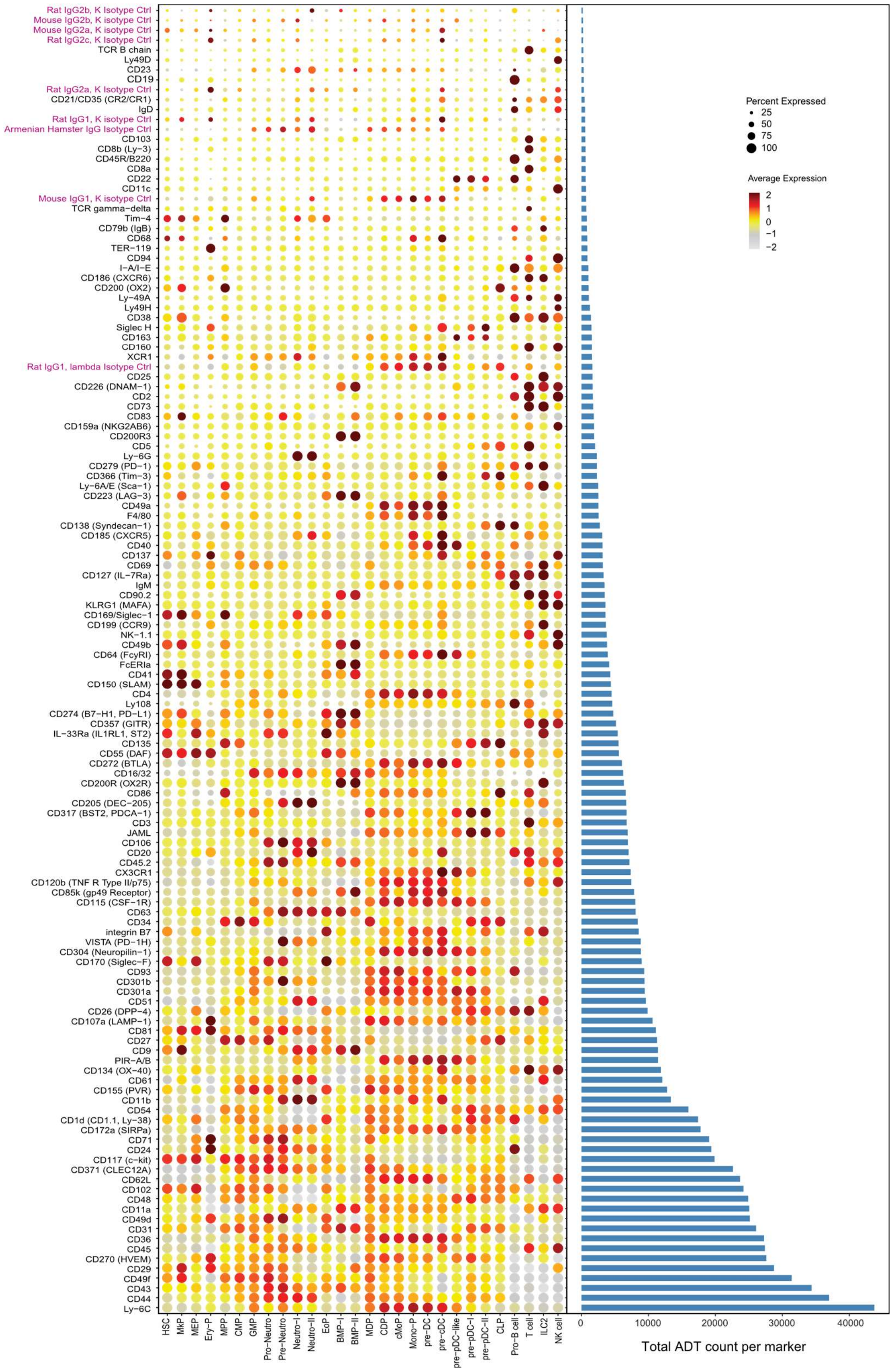


**Supplemental Figure 3:** Bead plot (left) showing the expression of all 123 cell-surface markers and isotypes across the BM hematopoietic progenitor stages. Mean expression is represented as colored-scale of relative intensities and the size of the circle represents the fraction of cells within that cluster having non-zero expression values. The bar plot (right) depicts the number of total reads obtained for each marker indicating the absolute expression level in the BM progenitor compartment.

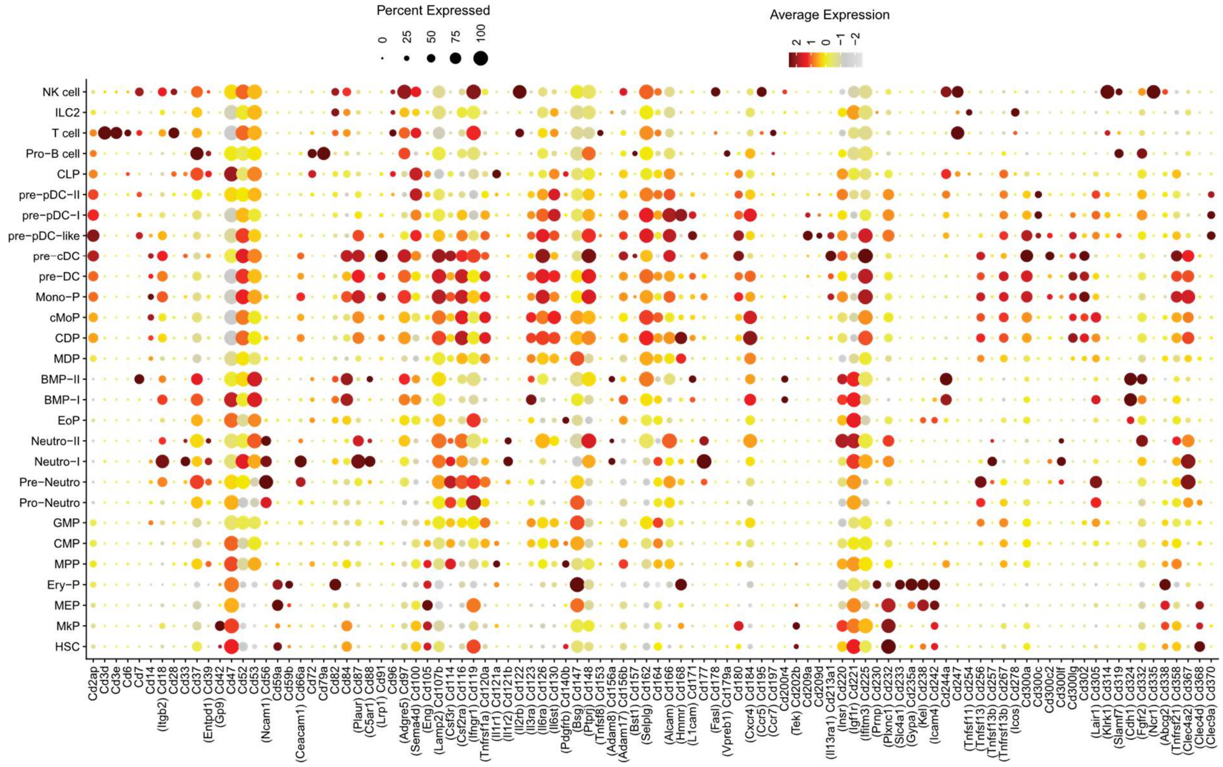


**Supplemental Figure 4:** Expression of selected non-ADT cell-surface CD markers at the transcript level across hematopoietic progenitor states. Markers are shown using standard CD nomenclature, with corresponding mm10 gene symbols indicated in brackets where they differ from the CD designation. Mean expression level is represented as colored-scale of relative intensities and the size of the circle represents the fraction of cells within that cluster having non-zero expression values.

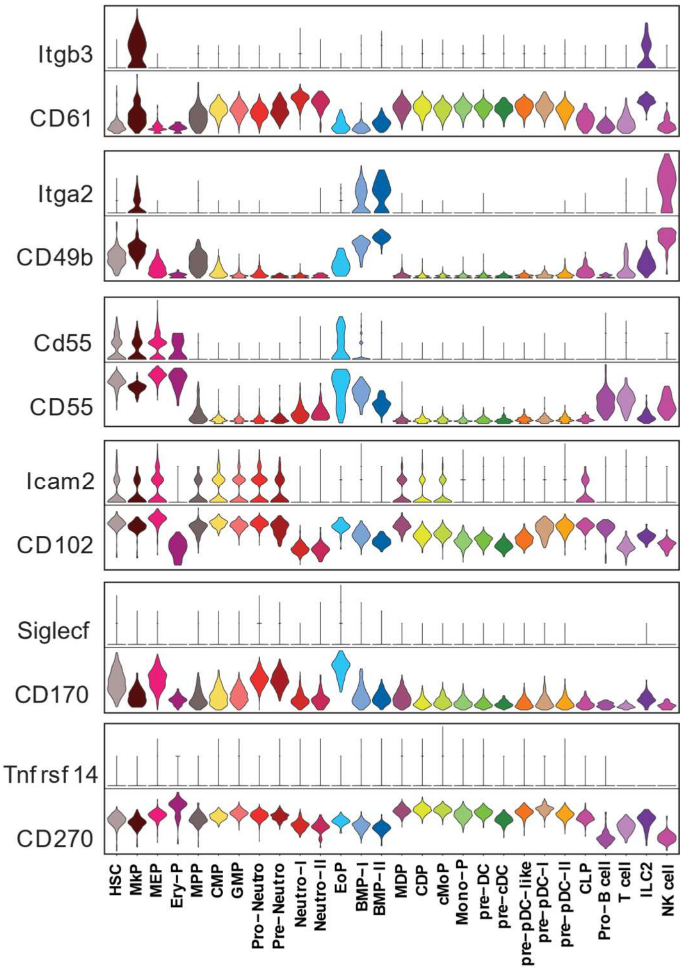


**Supplemental Figure 5:** Additional examples of protein versus transcript level expression discordance of cell surface markers. Violin plots comparing the transcript level expressions with the corresponding ADT-measured surface protein expressions across the progenitor stages. Each surface protein is grouped together with the corresponding transcript profile.

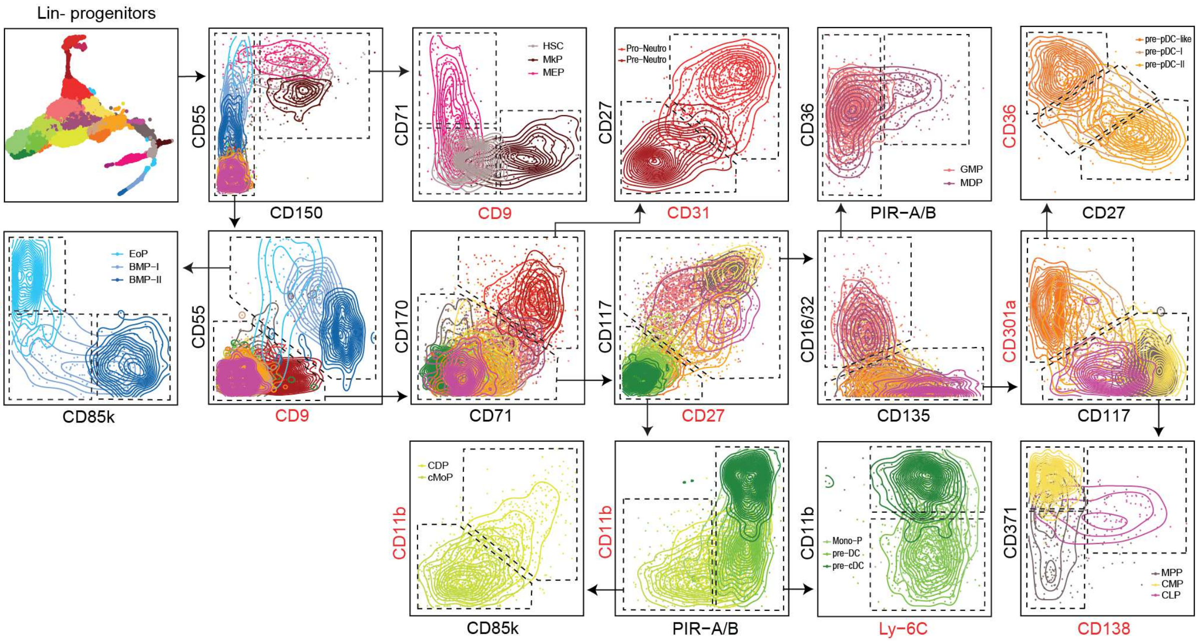


**Supplemental Figure 6:** Alternative surface markers for the gating strategy shown in Figure 7B. The replaced markers in the respective plots are color-coded as red. Dotted lines are used to show the proposed gates. Respective cluster annotations are mentioned in terminal gates. Lin = B220, CD3, CD4, CD8a, CD11b, CD11c, CD19, Ly-6G, MHC-II, NK1.1, and Ter119.

**Supplemental Tables**

Supplemental Table 1 FACS antibodies used for lineage positive cells depletion

Supplemental Table 2 Hash-tag antibodies used

Supplemental Table 3 CITE-seq antibodies in TotalSeq$^{TM}$-A Mouse Universal Cocktail

Supplemental Table 4 Spiked-in CITE-seq antibodies with titrated dilutions

**Supplemental Table 1:** FACS antibodies used for lineage positive cells depletion.

| Reactivity | Fluorophore | Marker | Clone | Dilution | Cat. No. | Company |
|---|---|---|---|---|---|---|
| Anti-mouse | FITC | CD3 | 145-2C11 | 1:100 | 553062 | BD Bioscience |
| Anti-mouse | FITC | CD4 | RM4-5 | 1:100 | 553047 | BD Bioscience |
| Anti-mouse | FITC | CD19 | 1D3 | 1:100 | 557398 | BD Bioscience |
| Anti-mouse | APC-Cy7 | CD11b | M1/70 | 1:100 | 557657 | BD Bioscience |
| Anti-mouse | APC | B220 | RA3-6B2 | 1:100 | 553092 | BD Bioscience |
| Anti-mouse | APC | NK1.1 | Pk136 | 1:100 | 550627 | BD Bioscience |
| Anti-mouse | PE | Ly-6G | 18A | 1:100 | 551461 | BD Bioscience |
| Anti-mouse | PE | Ter119 | TER-119 | 1:100 | 553673 | BD Bioscience |
| Anti-mouse | PE | I-A/I-E (MHCII) | N5/114.15.2 | 1:100 | 557000 | BD Bioscience |
| Anti-mouse | PE.Cy7 | CD8a | 53-6.7 | 1:100 | 552877 | BD Bioscience |
| Anti-mouse | PE.Cy7 | CD11c | HL3 | 1:100 | 558079 | BD Bioscience |

**Supplemental Table 2:** Hash-tag antibodies used.

| Reactivity | Marker | Clone | Barcode sequence | ID | Cat. No. | Company |
|---|---|---|---|---|---|---|
| Anti-mouse | Hashtag 1 | M1/42; 30-F11 | ACCCACCAGTAAGAC | A0301 | 155801 | Biolegend |
| Anti-mouse | Hashtag 2 | M1/42; 30-F11 | GGTCGAGAGCATTCA | A0302 | 155803 | Biolegend |
| Anti-mouse | Hashtag 3 | M1/42; 30-F11 | CTTGCCGCATGTCAT | A0303 | 155805 | Biolegend |
| Anti-mouse | Hashtag 4 | M1/42; 30-F11 | AAAGCATTCTTCACG | A0304 | 155807 | Biolegend |
| Anti-mouse | Hashtag 5 | M1/42; 30-F11 | CTTTGTCTTTGTGAG | A0305 | 155809 | Biolegend |
| Anti-mouse | Hashtag 6 | M1/42; 30-F11 | TATGCTGCCACGGTA | A0306 | 155811 | Biolegend |

**Supplemental Table 3:** CITE-seq antibodies in TotalSeq™-A Mouse Universal Cocktail.

| Reactivity | Surface marker | clone | ID | Cat. No. | Company |
|---|---|---|---|---|---|
| Anti-mouse | CD4 | RM4-5 | A0001 | 199901 | Biolegend |
| Anti-mouse | CD8a | 53-6.7 | A0002 | 199901 | Biolegend |
| Anti-mouse | CD366 (Tim-3) | RMT3-23 | A0003 | 199901 | Biolegend |
| Anti-mouse | CD279 (PD-1) | RMP1-30 | A0004 | 199901 | Biolegend |
| Anti-mouse | Ly-6C | HK1.4 | A0013 | 199901 | Biolegend |
| Anti-mo/hu | CD11b | M1/70 | A0014 | 199901 | Biolegend |
| Anti-mouse | Ly-6G | 1A8 | A0015 | 199901 | Biolegend |
| Anti-mo/hu | CD49f | GoH3 | A0070 | 199901 | Biolegend |
| Anti-mo/hu | CD44 | IM7 | A0073 | 199901 | Biolegend |
| Anti-mouse | CD54 | YN1/1.7.4 | A0074 | 199901 | Biolegend |
| Anti-mouse | CD90.2 | 30-H12 | A0075 | 199901 | Biolegend |
| Anti-mouse | CD73 | TY/11.8 | A0077 | 199901 | Biolegend |
| Anti-mouse | CD49d | R1-2 | A0078 | 199901 | Biolegend |
| Anti-mouse | CD200 (OX2) | OX-90 | A0079 | 199901 | Biolegend |
| Anti-mouse | CD19 | 6D5 | A0093 | 199901 | Biolegend |
| Anti-mouse | CD45 | 30-F11 | A0096 | 199901 | Biolegend |
| Anti-mouse | CD25 | PC61 | A0097 | 199901 | Biolegend |
| Anti-mo/hu | CD45R/B220 | RA3-6B2 | A0103 | 199901 | Biolegend |
| Anti-mouse | CD102 | 3C4 (MIC2/4) | A0104 | 199901 | Biolegend |
| Anti-mouse | CD115 (CSF-1R) | AFS98 | A0105 | 199901 | Biolegend |
| Anti-mouse | CD11c | N418 | A0106 | 199901 | Biolegend |
| Anti-mouse | CD21/CD35 | 7 E9 | A0107 | 199901 | Biolegend |

| | | | | | |
|---|---|---|---|---|---|
| Anti-mouse | CD23 | B3B4 | A0108 | 199901 | Biolegend |
| Anti-mouse | CD43 | S11 | A0110 | 199901 | Biolegend |
| Anti-mouse | CD5 | 53-7.3 | A0111 | 199901 | Biolegend |
| Anti-mouse | CD62L | MEL-14 | A0112 | 199901 | Biolegend |
| Anti-mouse | CD93 | AA4.1 | A0113 | 199901 | Biolegend |
| Anti-mouse | F4/80 | BM8 | A0114 | 199901 | Biolegend |
| Anti-mouse | FcεRIα | MAR-1 | A0115 | 199901 | Biolegend |
| Anti-mouse | I-A/I-E (MHC-II) | M5/114.15.2 | A0117 | 199901 | Biolegend |
| Anti-mouse | NK-1.1 | PK136 | A0118 | 199901 | Biolegend |
| Anti-mouse | Siglec H | 551 | A0119 | 199901 | Biolegend |
| Anti-mouse | TCR β chain | H57-597 | A0120 | 199901 | Biolegend |
| Anti-mouse | TCR γ/δ | GL3 | A0121 | 199901 | Biolegend |
| Anti-mouse | TER-119 | TER-119 | A0122 | 199901 | Biolegend |
| Anti-mouse | Ly-6A/E (Sca-1) | D7 | A0130 | 199901 | Biolegend |
| Anti-mouse | CD45.2 | 104 | A0157 | 199901 | Biolegend |
| Anti-mouse | CD3 | 17A2 | A0182 | 199901 | Biolegend |
| Anti-mouse | CD274 (B7-H1, PD-L1) | MIH6 | A0190 | 199901 | Biolegend |
| Anti-mo/hu/rat | CD27 | LG.3A10 | A0191 | 199901 | Biolegend |
| Anti-mouse | CD20 | SA275A11 | A0192 | 199901 | Biolegend |
| Anti-mouse | CD357 (GITR) | DTA-1 | A0193 | 199901 | Biolegend |
| Anti-mouse | CD137 | 17B5 | A0194 | 199901 | Biolegend |
| Anti-mouse | CD134 (OX-40) | OX-86 | A0195 | 199901 | Biolegend |
| Anti-mouse | CD69 | H1.2F3 | A0197 | 199901 | Biolegend |
| Anti-mouse | CD127 (IL-7Rα) | A7R34 | A0198 | 199901 | Biolegend |
| Anti-mouse | CD86 | GL-1 | A0200 | 199901 | Biolegend |
| Anti-mouse | CD103 | 2 E7 | A0201 | 199901 | Biolegend |
| Anti-mouse | CD64 (FcγRI) | X54-5/7.1 | A0202 | 199901 | Biolegend |
| Anti-mouse | CD150 (SLAM) | TC15-12F12.2 | A0203 | 199901 | Biolegend |
| Anti-mouse | CD24 | M1/69 | A0212 | 199901 | Biolegend |
| Anti-mo/hu | integrin β7 | FIB504 | A0214 | 199901 | Biolegend |
| Anti-mouse | CD106 | 429 (MVCAM.A) | A0226 | 199901 | Biolegend |
| Anti-mouse | CD8b (Ly-3) | YTS156.7.7 | A0230 | 199901 | Biolegend |
| Anti-mo/hu | KLRG1 (MAFA) | 2F1/KLRG1 | A0250 | 199901 | Biolegend |
| Anti-mouse | CD223 (LAG-3) | C9B7W | A0378 | 199901 | Biolegend |
| Anti-mouse | CD163 | S15049I | A0417 | 199901 | Biolegend |
| Anti-mouse | CD49b | HMα2 | A0421 | 199901 | Biolegend |
| Anti-mouse | CD172a (SIRPα) | P84 | A0422 | 199901 | Biolegend |
| Anti-mouse | CD48 | HM48-1 | A0429 | 199901 | Biolegend |
| Anti-mouse | CD170 (Siglec-F) | S17007L | A0431 | 199901 | Biolegend |
| Anti-mouse | CD169/Siglec-1 | 3D6.112 | A0440 | 199901 | Biolegend |
| Anti-mouse | CD71 | RI7217 | A0441 | 199901 | Biolegend |
| Anti-mouse | CD41 | MWReg30 | A0443 | 199901 | Biolegend |
| Anti-mouse | IgM | RMM-1 | A0450 | 199901 | Biolegend |
| Anti-mouse | CD301a | LOM-8.7 | A0551 | 199901 | Biolegend |
| Anti-mouse | CD304 (Neuropilin-1) | 3 E12 | A0552 | 199901 | Biolegend |
| Anti-mouse | CD36 | HM36 | A0555 | 199901 | Biolegend |
| Anti-mouse | CD38 | 90 | A0557 | 199901 | Biolegend |
| Anti-mouse | CD55 (DAF) | RIKO-3 | A0558 | 199901 | Biolegend |
| Anti-mouse | CD63 | NVG-2 | A0559 | 199901 | Biolegend |
| Anti-mouse | CD68 | FA-11 | A0560 | 199901 | Biolegend |
| Anti-mouse | CD79b (Igβ) | HM79-12 | A0561 | 199901 | Biolegend |
| Anti-mouse | CD83 | Michel-19 | A0562 | 199901 | Biolegend |
| Anti-mouse | CX3CR1 | SA011F11 | A0563 | 199901 | Biolegend |
| Anti-mouse | CD301b | URA-1 | A0566 | 199901 | Biolegend |
| Anti-mouse | Tim-4 | RMT4-54 | A0567 | 199901 | Biolegend |
| Anti-mo/rat | XCR1 | ZET | A0568 | 199901 | Biolegend |
| Anti-mo/rat | CD29 | HMβ1-1 | A0570 | 199901 | Biolegend |
| Anti-mouse | IgD | 11-26c.2a | A0571 | 199901 | Biolegend |
| Anti-mouse | CD11a | M17/4 | A0595 | 199901 | Biolegend |

| | | | | | |
|---|---|---|---|---|---|
| Anti-mouse | CD200R (OX2R) | OX-110 | A0807 | 199901 | Biolegend |
| Anti-mouse | CD200R3 | Ba13 | A0809 | 199901 | Biolegend |
| Anti-mouse | CD138 (Syndecan-1) | 281-2 | A0810 | 199901 | Biolegend |
| Anti-mouse | CD317 (BST2, PDCA-1) | 927 | A0811 | 199901 | Biolegend |
| Anti-mouse | CD9 | MZ3 | A0813 | 199901 | Biolegend |
| Anti-mouse | CD371 (CLEC12A) | 5D3/CLEC12A | A0825 | 199901 | Biolegend |
| Anti-mouse | CD22 | OX-97 | A0827 | 199901 | Biolegend |
| Anti-mouse | IL-33Rα (IL1RL1, ST2) | DIH9 | A0837 | 199901 | Biolegend |
| Anti-mouse | Ly49H | 3D10 | A0839 | 199901 | Biolegend |
| Anti-mouse | Ly49D | 4 E5 | A0841 | 199901 | Biolegend |
| Anti-mouse | Ly-49A | YE1/48.10.6 | A0842 | 199901 | Biolegend |
| Anti-mouse | CD185 (CXCR5) | L138D7 | A0846 | 199901 | Biolegend |
| Anti-mouse | CD49a | HMα1 | A0850 | 199901 | Biolegend |
| Anti-mouse | CD1d (CD1.1, Ly-38) | 1B1 | A0851 | 199901 | Biolegend |
| Anti-mouse | CD226 (DNAM-1) | 10 E5 | A0852 | 199901 | Biolegend |
| Anti-mouse | CD199 (CCR9) | CW-1.2 | A0854 | 199901 | Biolegend |
| Anti-mouse | JAML | 4 E10 | A0877 | 199901 | Biolegend |
| Anti-mouse | CD272 (BTLA) | 6A6 | A0881 | 199901 | Biolegend |
| Anti-mouse | PIR-A/B | 6C1 | A0882 | 199901 | Biolegend |
| Anti-mouse | CD26 (DPP-4) | H194-112 | A0883 | 199901 | Biolegend |
| Anti-mouse | CD270 (HVEM) | HMHV-1B18 | A0885 | 199901 | Biolegend |
| Anti-mouse | CD2 | RM2-5 | A0892 | 199901 | Biolegend |
| Anti-mouse | CD120b | TR75-89 | A0893 | 199901 | Biolegend |
| Anti-mouse | CD40 | 3/23 | A0903 | 199901 | Biolegend |
| Anti-mouse | CD31 | 390 | A0904 | 199901 | Biolegend |
| Anti-mouse | CD107a (LAMP-1) | 1D4B | A0905 | 199901 | Biolegend |
| Anti-mo/rat | CD61 | 2C9.G2 (HMβ3-1) | A0910 | 199901 | Biolegend |
| Anti-mouse | VISTA (PD-1H) | MIH63 | A0915 | 199901 | Biolegend |
| Anti-mouse | CD186 (CXCR6) | SA051D1 | A0926 | 199901 | Biolegend |
| Anti-mouse | CD159a (NKG2AB6) | 16A11 | A0927 | 199901 | Biolegend |
| Anti-mouse | Ly108 | 330-AJ | A0930 | 199901 | Biolegend |
| Anti-mouse | CD160 | 7H1 | A1006 | 199901 | Biolegend |
| Anti-mouse | CD85k (gp49 Receptor) | H1.1 | A1007 | 199901 | Biolegend |
| Anti-mouse | CD51 | RMV-7 | A1008 | 199901 | Biolegend |
| Anti-mouse | CD94 | 18d3 | A1009 | 199901 | Biolegend |
| Anti-mouse | CD205 (DEC-205) | NLDC-145 | A1010 | 199901 | Biolegend |
| Anti-mouse | CD155 (PVR) | TX56 | A1011 | 199901 | Biolegend |
| Anti-mo/rat | CD81 | Eat-2 | A1064 | 199901 | Biolegend |

**Supplemental Table 4:** Spiked-in CITE-seq antibodies with titrated dilutions.

| Reactivity | Surface marker | clone | Dilutions | ID | Cat. No. | Company |
|---|---|---|---|---|---|---|
| Anti-mouse | CD117 (c-kit) | 2B8 | 1:1,600 | A0012 | 105843 | Biolegend |
| Anti-mouse | CD135 | A2F10 | 1:100 | A0098 | 135316 | Biolegend |
| Anti-mouse | CD16/32 | 93 | 1:10,000 | A0109 | 101343 | Biolegend |
| Anti-mouse | CD34 | SA376A4 | 1:800 | A0823 | 152219 | Biolegend |
